# A maternally inherited Chromosomal Passenger Complex regulates germ plasm ribonucleoparticle aggregation in Zebrafish

**DOI:** 10.1101/2024.08.19.608573

**Authors:** Cara E. Moravec, Francisco Pelegri

**Author notes:** Correspondence to: Francisco Pelegri, Laboratory of Genetics, 2424 Genetics/Biotech, University of Wisconsin –Madison, 425-G Henry Mall, Madison WI 53706.

## Abstract

In zebrafish, the formation of primordial germ cells depends on the inheritance of a compartmentalized membrane-less subcellular structure containing a pool of maternally expressed germ plasm ribonucleoparticles (gpRNPs) and proteins. Interactions between cytoskeletal components and gpRNPs are crucial for the movement and collection of gpRNPs into the furrows during the first few cellular division of the early embryo. Previous work has identified *motley/ birc5b*, a maternally-expressed homolog of a known Chromosomal Passenger Complex (CPC) component, Survivin, as a linker between gpRNPs and microtubules during gpRNP aggregation. However, Survivin can also function independent of the CPC in other cellular contexts. Here we investigated whether a maternally inherited CPC is necessary for gpRNP aggregation. We identified *cdca9* as a maternally-expressed duplicated homolog of Borealin, another member of the CPC. Similar to *motley*, embryos from homozygous *cdca9* mutant females exhibit defects in chromosome segregation and cytokinesis during meiosis and mitosis, phenotypes associated with mutations in CPC members. Additionally, embryos lacking Cdca9 displayed decreased gpRNP aggregation prior to furrow formation in the early embryo, a phenotype indistinguishable from that observed in *motley* mutants. As previously shown for Birc5b, Cdca9 and other CPC components INCENP and Aurora B kinase colocalize at the tips of astral microtubules as gpRNPs are transported to the forming furrow. Unexpectedly, Birc5b, but not other CPC components, accumulates within the growing gpRNP aggregate prior to and during furrow formation. The association of Birc5b with germ plasm masses continues during their asymmetric segregation in the cleavage stages, ceasing only when gpRNPs undergo cytoplasmic dispersal during gastrulation. Our studies reveal a role for a non-conventional, maternally-inherited CPC for spindle and furrow formation, and, unexpectedly, gpRNP aggregation during early development. Additionally, we find that Birc5b, but not other CPC proteins, remains a component of zebrafish germ plasm during and after its aggregation.

**Author Summary:** Maternal products are necessary for early development across species, and the removal of these products from the embryo can cause developmental defects or death. The zebrafish has been widely used to discover the role of maternal products during early development. Using zebrafish, we discovered that a mutation in a maternal-specific duplicated *borealin* gene not only affects early development but also the aggregation of germ cell determinants. We also find that this duplicated Borealin interacts within a specialized Chromosomal Passenger Complex, a complex that traditionally regulates multiple steps of cellular division. This specialized Chromosomal Passenger Complex acts as a linker between germ cell determinants and the cytoskeleton during early development. These results highlight a unique role for the Chromosomal Passenger Complex outside of cellular division during early development. Further, these findings underscore the intricate mechanisms by which gene duplications contribute to the regulation of early developmental processes, providing valuable insight into the molecular events of embryogenesis.

## Introduction

In animal species, embryonic development depends on a pool of maternal products for all cellular functions from fertilization until zygotic genome activation, including the regulation of cellular division and cell fate determination (1–3). One of the first cell fate decisions that occurs in early development is the specification of the primordial germ cells (PGCs). In zebrafish, PGC specification requires the aggregation of maternally supplied RNAs and proteins into a biomolecular condensate, known as germ plasm (4–7). Zebrafish germ plasm is maternally inherited as germ plasm ribonucleoparticles (gpRNPs) that aggregate to form four masses at the furrow distal ends (7–14). These distal masses ingress into a cell and will later in development confer the germ cell state (10,15–17).

The transport of RNPs to and within the furrow relies on the coordination of the cytoskeleton, specifically involving microtubule and actin networks (7,8,11,12,14,18–20). Prior to furrow formation, germ plasm RNPs aggregate at the tips of the astral microtubules (12). As growing astral microtubules extend outwardly, they are thought to move gpRNPs towards the edge of the blastodisc (12) with this movement, coupled to particle-particle interactions, promoting gpRNPs aggregation. Additionally, cortical F-actin rings appear to modulate the outward movement of the gpRNPs, facilitating the recruitment of germ plasm RNPs to the forming furrows (19).

Once the gpRNPs are recruited to the furrow, they become enriched at their distal ends (8,10,13,14,19,19) in a process involving dynamic cytoskeletal changes during furrow maturation. At the furrow and in addition to F-actin filaments, the microtubule apparatus forms a furrow microtubule array (FMA), characterized by a ladder-like arrangement of microtubule bundles parallel to each other and perpendicular to the furrow (21,22). Similar to F-actin, this microtubule array is required for cytokinesis, with the FMA thought to function in the secretion of membrane vesicles to the furrow site (21,22). As furrow maturation progresses, the FMA also undergoes reorganization, accumulating bundling microtubule ends at distal furrow regions (13,22). During this process, FMA microtubule bundled tips are associated with gpRNP aggregates recruited at the furrow, such that they coordinately accumulate at furrow distal ends resulting in microtubule-associated, compact germ plasm masses in these locations (20). In addition to its role in membrane ingression during cytokinesis, furrow F-actin exhibits wave-like repetitive contractions required for the compaction of gpRNPs to the furrow’s distal ends (9,19). Birc5b, a duplicated homolog of Survivin (Birc5), is a maternally expressed protein that is required both for cytokinesis and gpRNPs aggregation in the early embryo (12). One major mitotic function in the cell for Survivin/Birc5 is as a member of the Chromosomal Passenger Complex (CPC) (23), which has regulatory roles at multiple steps of the cell cycle including the localization of chromosomes to the metaphase plate, cohesion protection, biorientation error correction, and membrane ingression during cytokinesis (24–27). The CPC comprises four members, the serine/threonine kinase, Aurora Kinase B and three localization factors, Survivin, Borealin and INCENP, that interact via an interdependent coiled-coil triple helical bundle responsible for the localization of the CPC to chromosomes and microtubules (Adams et al., 2000; Adams et al., 2001; Gassmann et al., 2004; Jeyaprakash et al., 2007a; Jeyaprakash et al., 2011; Klein et al., 2006; Romano et al., 2003; Sampath et al., 2004; Vader et al., 2006; Wheatley et al., 2001). Apart from its involvement in the CPC, Survivin/Birc5 plays diverse cellular roles encompassing functions such as cell survival, the regulation of focal adhesions, and epithelial-mesenchymal transition (28–31).

Here, we asked whether Birc5b acts within a maternally inherited CPC in the regulation of gpRNP aggregation in the early zebrafish embryo cortex. We identified *cdca9,* a maternal-specific duplicated gene for *borealin/cdca8* and find that, similar to *motley*/*birc5b* mutants, embryos lacking maternal Cdca9 function exhibit failed cytokinesis and defects in gpRNP aggregation. The localization of other CPC members, INCENP and Aurora Kinase B, suggest that all four proteins form part of a CPC that acts as a linker between the microtubules ends and aggregating gpRNPs. Unexpectedly, Birc5b, but not other CPC components, accumulates in the growing gpRNP aggregate, an association that continues during the recruitment and compaction of gpRNPs at the furrow to form the germ plasm until gpRNPs undergo cytoplasmic dispersal during mid-gastrulation. Our studies show that a maternally-inherited CPC is a crucial interactor between the cytoskeleton and aggregating gpRNPs. Additionally, we find that Birc5b, but not the other CPC members, is a core factor associated with zebrafish germ plasm during its aggregation and segregation in early embryonic development.

## Results

### Identification of a maternally expressed zebrafish *borealin*

Our previous studies showed that Birc5b, a gene duplicate for Survivin with maternal-specific expression, is required for cytokinesis and gpRNP aggregation in the early embryo. On the other hand, the duplicate homolog, *birc5a*, is closely related to Survivin/Birc5 in other lineages and is expressed throughout development, suggesting a more general involvement in fundamental cellular functions throughout the lifetime of the organism. These findings suggested that the duplication of an ancestral *birc5* gene enabled the subfunctionalization of the *birc5b* gene copy for early developmental functions (12). If Birc5b functions within a specialized maternally-expressed Chromosomal Passenger Complex during the egg-to-embryo transition, we hypothesized that other members of the complex may have undergone a similar gene duplication event and subfunctionalization process. We thus examined the zebrafish genome and transcriptome for gene duplicates with maternal-specific gene expression for *borealin*, *aurora kinase b*, and *incenp*.

Using BLAST and a zebrafish mRNA baseline expression database (32), we determined that *borealin* has two copies with different expression patterns, *cdca8* and *cdac9* (Figures 1A,C)(33). In the zebrafish embryo, *cdca8* is present at low levels during cleavage stages, and increased levels are observed starting at the 128-cell stage. On the other hand, *cdca9* is found at high levels in the one-cell embryo, with the transcripts becoming undetectable prior to the segmentation stages (Figure 1A). We confirmed the *borealin* gene homologs expression via RT-PCR (Figure 1B). This pattern is similar to the zebrafish *birc5a*/*birc5b* expression with one homolog, either *birc5a* or *cdca8* expressed throughout development and a second copy, *birc5b* or *cdca9*, expressed only maternally (Figure 1A)(12). In contrast, *aurora kinase b* and *incenp* only have one gene copy that is expressed throughout development (Figure 1A).

**Figure 1:**
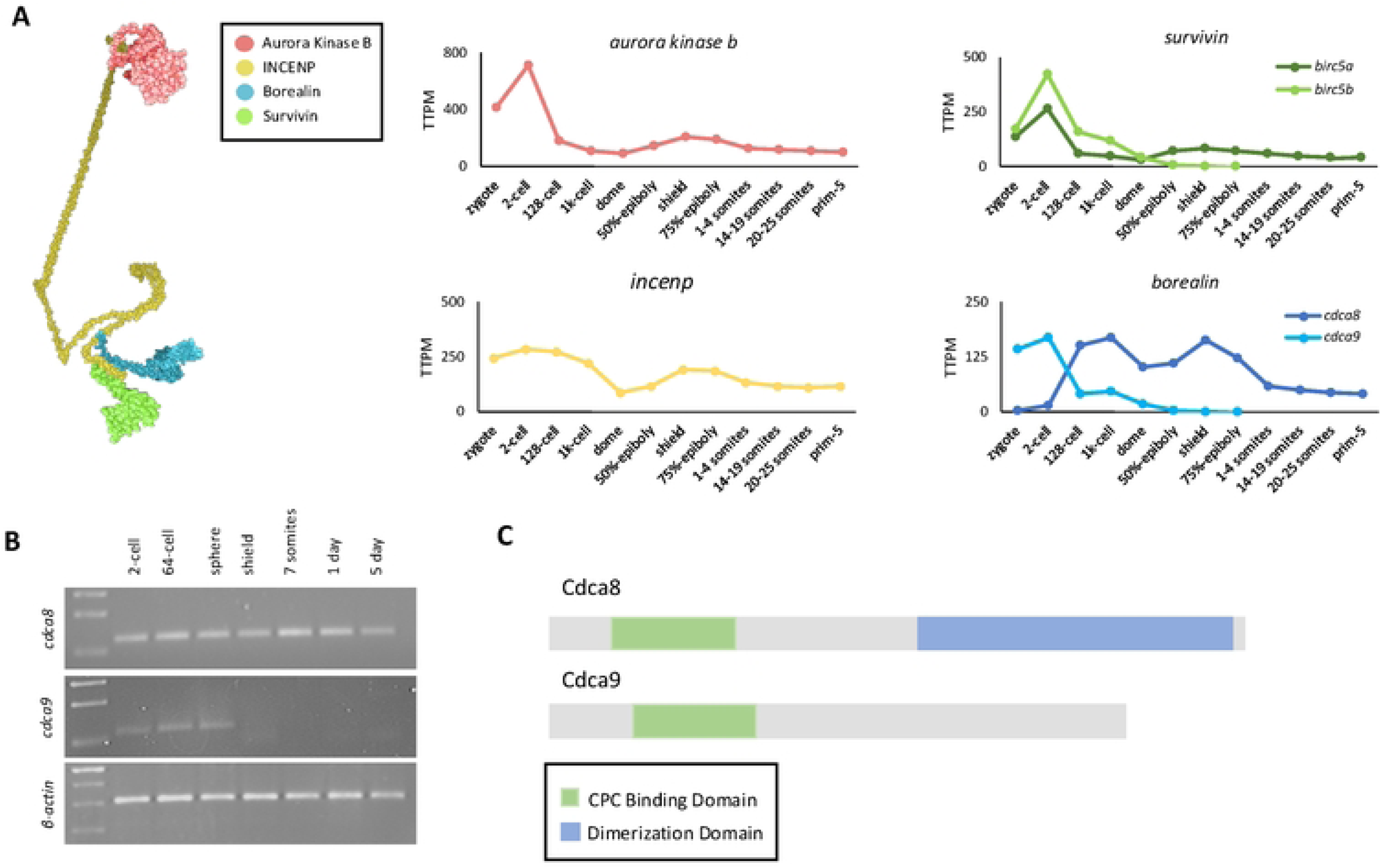
Identification of a maternally expressed *borealin* gene. A) On the left, a diagram showing relative placement and interactions of CPC components within the complex. Expression levels of CPC members during early development from zygote to Prim-5 (24hpf) stage. Both *survivin* and *barealin* have two gene copies, with one of the gene copies expressed only maternally during early development and the other at all stages. B) RT-PCR confirming mRNA expression levels of *cdca8* and *cdca9* up to 5 days after fertilization. C) Protein domain diagrams of Cdca8 and Cdca9 showing the known motifs of the two paralogs predicted by Pfam.

The maternal-specific expression patterns of Birc5b and Cdca9 combined with the known direct interaction of Borealin and Survivin within the conventional CPC (25–27), suggested Cdca9 could act as a candidate partner for Birc5b in a maternally-inherited CPC. To investigate Cdca9 role in early development, we targeted *cdca9* with CRISPR-Cas9 to generate a null allele. To prevent the formation of a truncated Cdca9 protein that would be able to interact with a CPC in a dominant negative fashion, we used guide RNAs targeted to the CPC interacting domain, located in the N-terminus and necessary for Borealin to interact with Survivin and INCENP (25). Subsequently, two unique alleles, *cdca9^UW102^* and *cdca9^UW103^* were isolated, which harbored deletions of 5 bp and 13 bp, respectively. Both alleles are predicted to make proteins that are less than 52 amino acids long compared to the full length 255 amino acids Cdca9 protein (Figure 2A-D) and generate products shorter than the minimal region of 109 amino acids in the CPC interacting domain (25). Thus, these mutant variants likely function as null alleles.

**Figure 2:**
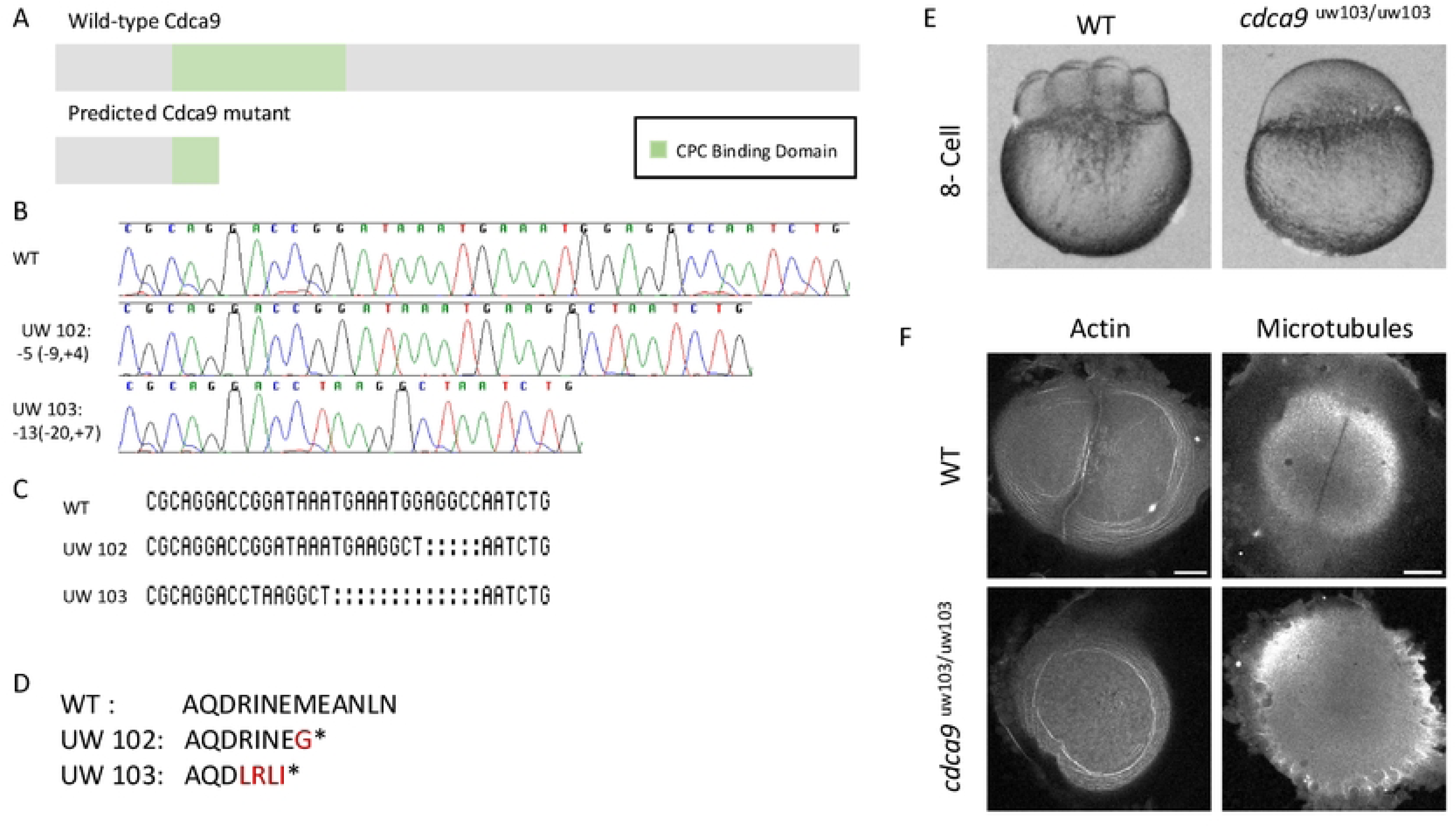
Cdca9 is necessary for cytokinesis. A) Diagrams highlighting the known conserved motif (CPC Binding Domain) in wild-type Cdca9 and the predicted truncated product encoded by *cdca9* CRISPR-Cas9 alleles. B,C) Sequence chromatograms (B) and alignment (C) for wild-type, *cdca9^UW102/UW102^*, and *cdca9^UW103/UW103^* alleles over the target site. (D) Predicted amino acid changes are highlighted in red, with premature stop codons indicated by asterisks in both *cdca9* CRISPR-Cas9 alleles. E) Live wild-type and cdca9 mutant embryos at the 8-cell stage (1 hour 15 minutes post fertilization. *cdca9* mutant embryos fail to form furrows. F) Labeling for microtubules and actin in fixed two-cell wild-type and *cdca9^UW103/UW103^* embryos showing the lack of a furrow in the *cdca9* mutant. Micrographs are 2D maximum projections of confocal Z-stacks, scale bars are 100 μm.

Homozygous *cdca9* mutant are viable, and mature into adults in the expected mendelian ratios, with embryos from adult mutant females exhibiting 100% embryonic lethality even when crossed to wild-type males (Figure 2E). For simplicity, we refer to embryos from homozygous mutant females as *cdca9* mutant embryos. Embryos generated from *cdca9^uw102/uw103^* transheterozygous females also showed 100% embryonic lethality, confirming that the phenotype is caused by depletion of wild-type Cdca9 from the early embryo (Supplementary Figure 1) and not an off-target mutation. Embryos from *cdca9* mutant males are morphologically normal (data not shown), consistent with a maternal effect.

### Cdca9 is required for cytokinesis in the early embryo

*cdca9* mutant embryos undergo fertilization and egg activation, but not cell cleavage (Figure 2E). In fixed embryos as assessed by β-catenin and DAPI at 4 hours post-fertilization, wild-type controls have established cell boundaries and discrete nuclei. In contrast, *cdca9* mutants lack cell boundaries and contain irregular-sized nuclei (Supplementary Figure 1A,B). These irregular nuclei could arise from destabilizing the mitotic spindle and/or a lack of formation of cell membranes during cell division, both phenotypes known to depend on CPC activity (12,33–36).

Labeling for actin shows that the *cdca9* mutants fail to establish a furrow. Wild-type embryos exhibit two distinct cellular structures: actin bundles along the furrow akin to the cytokinetic contractile ring, and actin arcs oriented circumferentially at the edge of the blastodisc. Time-matched *cdca9* mutant embryos fail to form an actin filament bundle corresponding to the contractile ring at the furrow, although blastodisc circumferential actin arcs appear to form normally (Figure 2F).

As the furrow develops in wild-type embryos, microtubules align parallel to each other and perpendicular to the furrow to create the Furrow Microtubule Array (FMA) (21,22). At the same time, an area free of microtubules, the microtubule exclusion zone, develops along the site of the furrow. *cdca9* mutant embryos fail to establish a microtubule exclusion zone and lack an FMA, with astral microtubules from both poles instead continuing to grow into a disorganized mesh-like structure that spans the site of furrow (Figure 2F). Thus, Cdca9 is necessary for proper formation and organization of the furrow cytoskeleton in the early embryo. The phenotypes observed in *cdca9* mutants are similar to those reported in *motley* mutants (12).

### *cdca9* is required for meiosis II

As with many other animal species, the mature zebrafish oocyte is arrested in metaphase of meiosis II. Upon egg activation, meiosis II proceeds, resulting in the extrusion of the second polar body. It has been previously reported that CPC components are required for meiosis in various species (37–41), including Birc5b in zebrafish (12). To evaluate the role of Cdca9 in meiosis II, we assessed the completion of this process in water-activated *cdca9* mutants.

In wild type, sister chromatids begin to segregate to the meiotic spindle poles immediately after egg activation (5 mpa, Fig. 3A, N=5). Subsequently, at 10 mpa, microtubules in the spindle midzone reorganize into a meiotic spindle between the two sets of segregating sister chromatids, with a midzone corresponding to the meiotic midbody at the site of abscission (Fig. 3B, gold arrow, N=6). By 20 mpa, one set of chromatids remains in the egg to form a female pronucleus, while the other chromatid set is extruded to generate the polar body, with the spindle remnant forming a midbody-like structure that is asymmetrically located, closer to the forming polar body. During this process, the female pronucleus appears to undergo decondensation and the polar body condensation, as assessed by size and intensity of labeling for a DNA dye (Fig. 3C, N=4).

**Figure 3:**
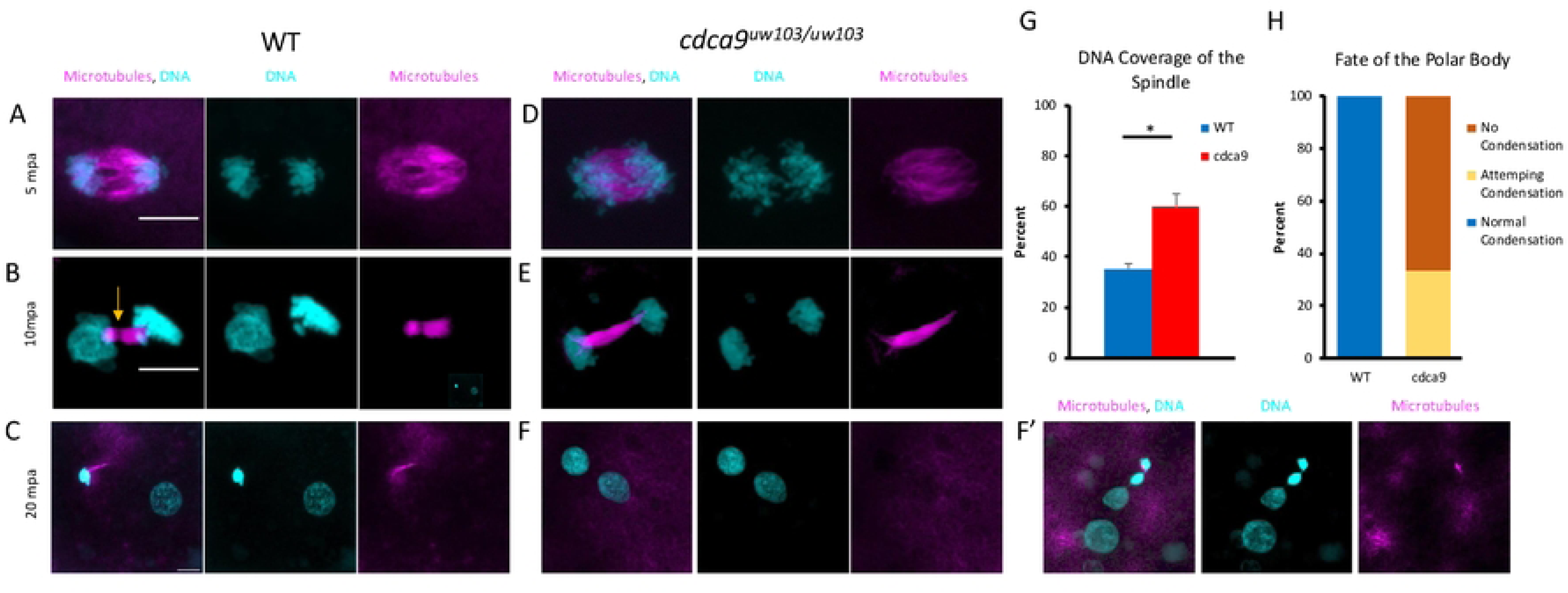
*cdca9* mutant eggs fail to complete meiosis II. Progression of meiosis II in water-activated wild-type (A-C) and *cdca9* mutant eggs (D-F). A) At 5 mpa, sister chromatids align at meiotic spindle poles during anaphase. B) At 10 mpa, the meiotic spindle bundles into a nascent midbody (arrow, notice gap in the microtubule structure) between one set of decondensing chromatids (the future female pronucleus) and one set of condensing chromatids (the future polar body). C) By 20mpa, the larger decondensed female pronucleus and the smaller condensed polar body have separated, with the meiotic midbody attached to the polar body (arrowhead). D) At 5 mpa in *cdca9* mutant eggs, sister chromatids appear more spread along the meiotic spindle. E) By 10 mpa, sister chromatid sets cannot be distinguished as either pronucleus or polar body and no meiotic midbody is seen in the *cdcda9* mutants. F) At 20 mpa, sister chromatid sets appear separate but exhibit similar degrees of condensation, suggesting that the 2^nd^ polar body did not form. In a fraction of cases (F’), DNA presumably corresponding to the polar body complement appears as a string of masses with various degrees of condensation, suggesting partially succesfully attempts at polar body formation. G) Percent coverage of chromosome spread over the mitotic spindle, showing that chromosomes are more dispersed in *cdca9* mutants. H) Degree of DNA condensation in the presumed polar body, as assessed by apparent nuclear mass volume and labeling intensity, consistent with lack of polar body extrusion due to defects in meiotic cytokinesis. All micrographs are 2D maximum projections of confocal Z-stacks, scale bars are 10 μm.

In water-activated eggs from *cdca9* mutant females, the meiotic spindle appears to initiate anaphase normally (Fig. 3D, N=14). However, the meiotic spindle length in *cdca9* mutants is significantly shorter than wild type (*cdca9*: 11.82 μm, N=10; WT: 14.98 μm, N=10, P-value=0.015). Additionally, segregating chromatids appear more widely distributed along the spindle in *cdca9* mutants when compared to their greater restriction at the spindle poles in wild type (Fig. 3G, *cdca9*: 60% chromosome coverage along spindle length, N=10; WT: 35%, N=10, P-value=0.0005), suggesting defects in chromosome segregation to the spindle poles.

At 10 mpa in *cdca9* mutants, the meiotic spindle lacks the gap in microtubule labeling corresponding to the meiotic midbody normally observable in wild-type (Fig. 3E, N=5). At 20mpa, *cdca9* mutants exhibit defects consistent with partial or failed extrusion of the polar body from the activated egg, such as a reduction or lack of condensation of the DNA mass corresponding to the polar body nucleus (N=6) (Figure 3F,F’,H). These meiotic phenotypes are similar to those described in water-activated eggs from *motley* mutant females (12).

### Cdca9 is required for germ plasm RNP multimerization

Germ plasm RNPs begin to aggregate at the cortex of the blastodisc even prior their recruitment to the forming furrows corresponding to the first embryonic cell division. During gpRNP aggregation, radially growing astral microtubules move the gpRNPs, promoting the formation of increasingly larger aggregates. Previous research has shown that, Birc5b acts as a linker between astral microtubules and is required to form high-order gpRNP aggregates at the cortex (Nair et al., 2013a).

To assess if *cdca9* has a role in gpRNP aggregation, we visualized gpRNP aggregation using an antibody against the human phosphorylated non-muscle myosin (NMII-p), a marker for gpRNPs in zebrafish (12). During their aggregation and recruitment to the furrow, gpRNPs maintain their distinct borders even when found in higher-order aggregates, allowing for semi-quantitative analysis of RNP number within the aggregates. In wild-type, the combined action of growing astral microtubules and particle-particle interactions is thought to result in gpRNP aggregation, as observed by the accumulation of clusters of various sizes, mid-size aggregates (2-6 gpRNPs/cluster) and larger aggregates (≥7 gpRNPs/cluster), in addition to the presence of remaining single gpRNPs that have not yet coalesced with neighboring particles.

Within this distribution at 40 mpf in wild-type during furrow formation, we observed 36.9% clusters containing at least 2 gpRNPs, including 2.7% containing 7 or more gpRNPs (N=1539, ROIs=39) (Figure 4 A,E,F, Table 2). In *cdca9* mutants, the fraction corresponding to clusters containing at least 2 gpRNPs was significantly reduced to 17.8 % (0.28% with ≥ 7 gpRNPs) (N=1064, embryos, ROIs=39) (Figure 4 B,E,F). Conversely, the single RNP category was significantly greater in *cdca9* mutants than wild type (*cdca9*: 82.0%, WT: 63.1%) (Figure 4 D, Table 2)). This reduction in aggregation is similar to that observed in *motley/birc5b* mutants (75.6% single RNP, 23.8 % 2-6 RNPs, 0.57% ≥7 RNPs) (Figure 4C,D-F, Table 2, N=1059, ROIs=39), as previously reported (12). Thus, *cdca9*, similar to *motley/birc5b*, is required for gpRNP aggregation in the cortex of the early embryo.

**Figure 4.**
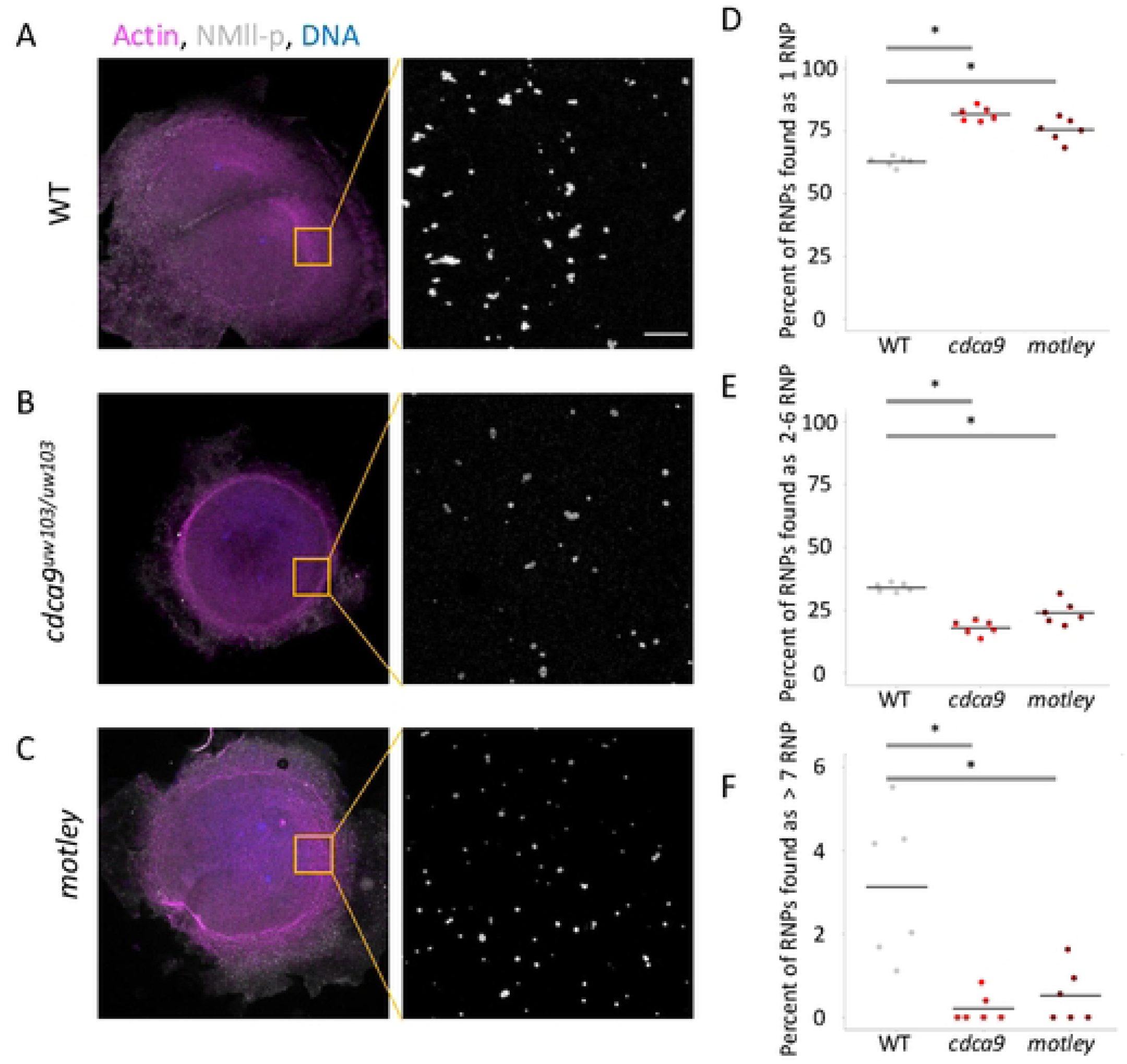
*cdca9* mutant fails to effectively multimerize gpRNP. A-C) Images of gpRNPs at 40 mpf forming aggregates at the cortex prior to their recruitment at the furrow. A) Wild-type embryos show gpRNP multimerization resulting in larger aggregates. B,C) Both *cdca9* and *motley* mutants exhibit a reduction in the number of larger gpRNP aggregates compared to wild-type embryos. D-F) Quantification of aggregation according to size class, showing a significant increase in single RNPs and a decrease in the frequency of larger aggregates. Asterisks indicate statistical significance. All micrographs are 2D maximum projections of confocal Z-stacks of animal views of blastodisc cortex, scale bars are 10 μm.

**Table 2:** Distribution of RNPs aggregate categories in *cdca9* and *motley* mutants compared to wild-type control embryos.

| Genotype | 1 RNP | 2-6 RNP | >7 RNP |
| --- | --- | --- | --- |
| WT | 63.0% | 34.5% | 2.7% |
| *cdca9* | 82.0% | 17.5% | 0.28% |
| P-value compared to WT | < 0.0001 | < 0.0001 | 0.0025 |
| *motley/birc5b* | 75.6% | 23.8% | 0.57% |
| P-value compared to WT | < 0.0001 | 0.0005 | 0.0067 |

### Birc5b, not Cdca9, accumulates within germ plasm RNP aggregates in the cortex

We asked if Cdca9 protein, like Birc5b, is localized to gpRNPs aggregates, labeled with NMII-p, in the early embryo. In the blastodisc of a wild-type embryo at 40 mpf, we observed that Cdca9 protein is present at the tip of microtubules in distinct puncta (Figure 5A). Notably, the bulk of Cdca9, does not localize to gpRNP aggregates, except in cases where microtubule and gpRNPs appear to be associated. In such cases, intensity profile tracing along the microtubule into the gpRNP aggregates revealed that Cdca9 is positioned between the microtubules and gpRNPs (Figure 5 B,C, N=5, ROIs=12). In contrast, in similar labelings to detect Birc5b protein in the blastodisc of the embryo, the majority of Birc5b and NMII-p fully colocalizes (Figure 5 D-F, N= 4, ROIs=12). Moreover, similar intensity profile tracing shows that Birc5b maxima are closer than the Cdca9 maxima to NMII-p maxima (P-value < 0.0001, Cdca9-0.67 μm, Birc5b-0.11 μm) (Figure 5 G).

**Figure 5:**
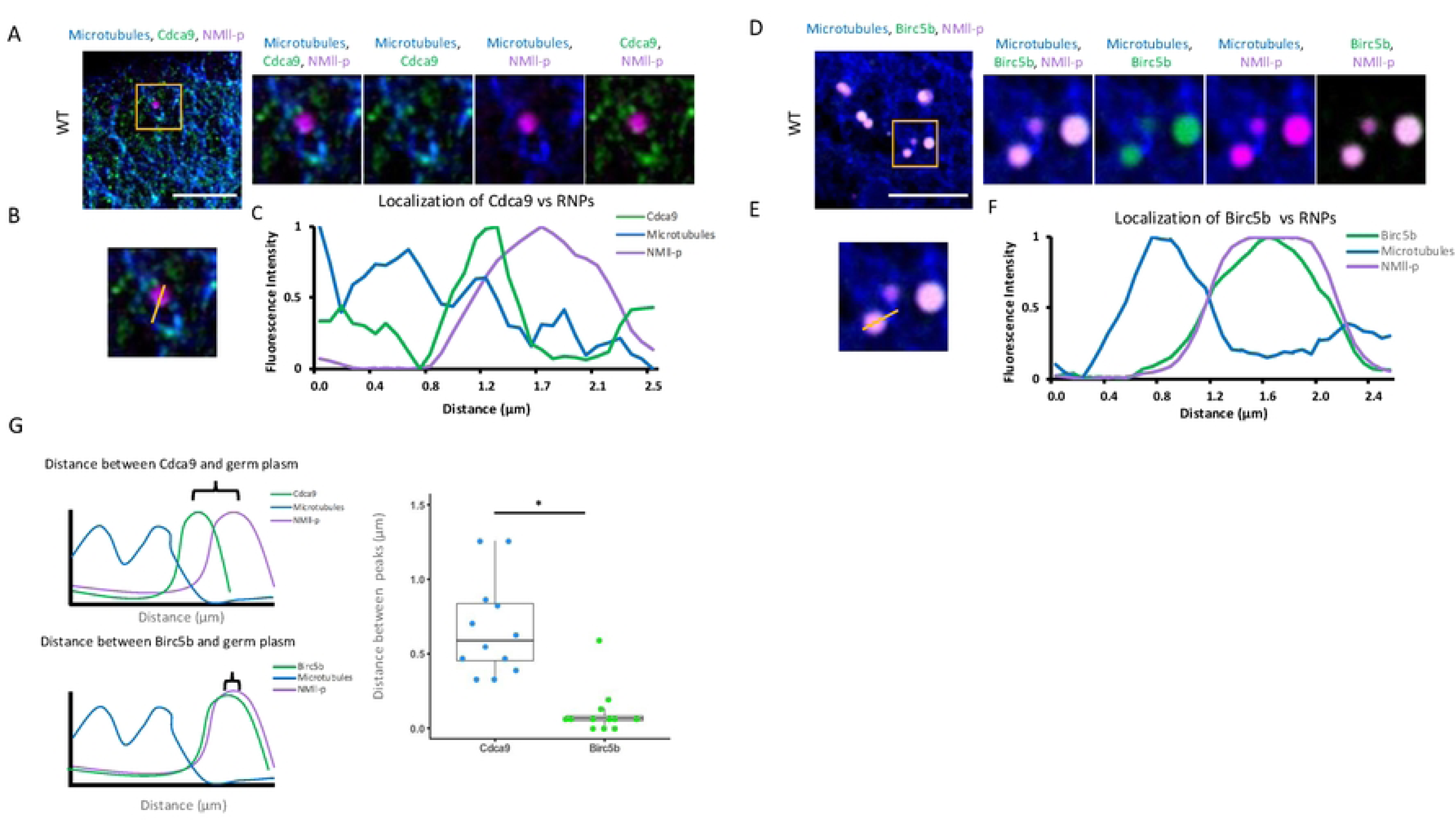
A majority of Cdca9 protein is localized to the tip of astral microtubules. A) Immunolabeling of microtubules, Cdca9 and NMll-p in the cortex of a wild type one-cell embryo. B,C) Line scan across the cortex shows that the majority of Cdca9 is located between the microtubules and NMll-p. D) Immunolabeling of microtubules, Birc5b and NMll-p in the cortex of a wild-type one-cell embryo. E,F) Line scan across the cortex shows that the majority of Birc5b colocalizes with NMll-p. G) Quantification of the distance between the two maxima between either Cdca9 or Birc5b and NMll-p, showing that maxima for Cdca9 are significantly farther away from the NMll-p maxima than Birc5b. All micrographs are 2D maximum projections of confocal Z-stacks, scale bars are 10 μm.

Thus, despite both mutants showing reduced gpRNP aggregation, Birc5b and Cdca9 show different localization patterns in the cortex of the one-cell embryo, with a majority of Birc5b (but not Cdca9) colocalizing with growing gpRNP aggregates. In cases where microtubules are associated with gpRNPs aggregates, Cdca9 is positioned between microtubules and gpRNP, whereas the bulk of Birc5b fully colocalizes with the gpRNP aggregate.

### Cdca9 is found in a maternally-expressed CPC on the tips of the astral microtubules

To assess whether Cdca9 could be acting within a maternal-expressed CPC in the early embryo, we examined the protein localization of the other three CPC members to astral microtubule tips. In wild-type embryos, we observed distinct puncta of INCENP, Aurora Kinase B, and Birc5b on the tips of astral microtubules, colocalizing with Cdca9 (Figure 6). In double labeling experiments involving Cdca9 and each of the other CPC components, fluorescent intensity scans in the Z-axis direction (into the cortex) of these puncta shows colocalization at astral microtubules ends for Cdca9 and INCENP (Figure 6 A-C, Pearson coefficient= 0.65, ROIs= 30), Aurora Kinase B (Figure 6 E-G, Pearson coefficient= 0.84, ROIs= 30) and Birc5b (Figure 6 I-K, Pearson coefficient= 0.48, ROIs= 30). As expected, a 90-degree rotated control showed no colocalization (Pearson coefficient= −0.04, Supplementary Figure 2, ROIs= 10). The data supports the idea that all CPC components colocalize at the tips of astral microtubules.

**Figure 6:**
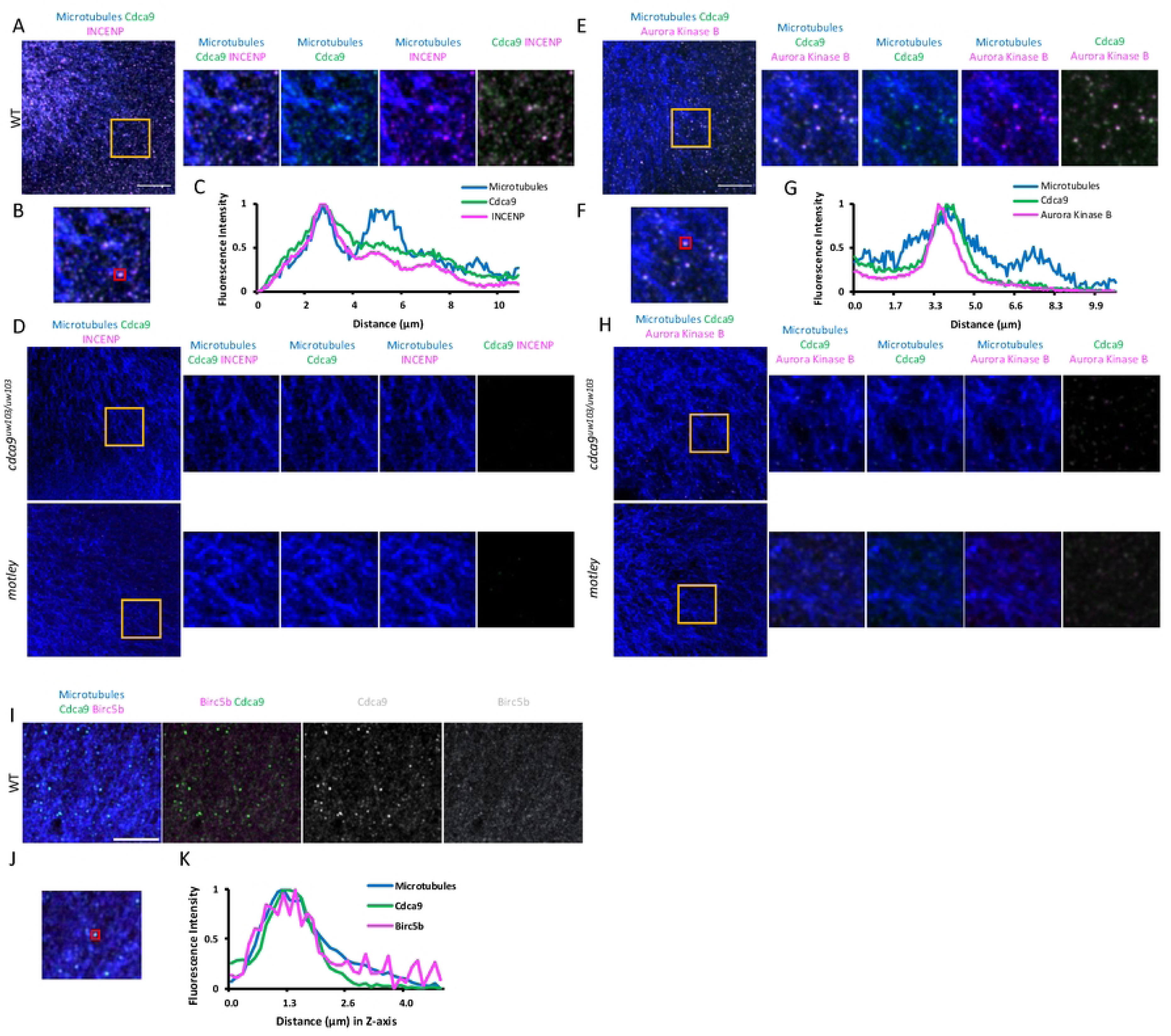
A maternally expressed CPC is found on the tips of astral microtubules. A-H) Immunolabeling of Cdca9 with either INCENP (A-D) or Aurora Kinase B (E-H). In wild-type Cdca9 colocalizes with INCENP (A-C) and Aurora Kinase B (E-G) at the tips of astral microtubules. The gold boxes in (A,E) shows the magnified views in images to the right and in (B,F), respectively. Punctae (red boxes) were analyzed using relative fluorescence intensity analysis in the z-direction (into the cortex), with the traces confirming colocalization of Cdca9 with both INCENP (C) and Aurora Kinase B (G) to a subset of microtubules. In *cdca9* or *motley* mutants, (D,H) no punctae are observed for either Cdca9, INCENP or Aurora Kinase B using the same imaging conditions. I-K) Immunolabeling of astral microtubules in a one-cell wild-type embryo showing colocation of Birc5b and Cdca9 at the tips of astral microtubules. The imaged area corresponds to a cortical region that does not include the furrow containing gpRNPs with larger Birc5b aggregates to allow visualization of lower levels of Birc5b at microtubule tips. K) Punctae (red boxes in J) were analyzed as in (C,G) confirming colocalization of Birc5b and Cdca9 at microtubule tips. All micrographs are 2D maximum projections of confocal Z-stacks, scale bars are 10 μm.

Prior research has shown that a stable and functional CPC requires all proteins and if any one of the proteins is missing, the complex dissociates (25,27,33,35,36). To further assess whether the hypothesized maternally expressed CPC behaves similarly to a canonical CPC, we performed immunolabeling analysis to determine the colocalization of CPC components to astral microtubules in *motley* and *cdca9* mutants. No distinct punctae of Cdca9, INCENP, or Aurora B kinase were observed on astral microtubules in the *cdca9* or *motley* mutants (Figure 6D,H). Thus, the colocalization of CPC proteins to microtubule tips are interdependent, consistent with these proteins forming a maternal CPC.

We used co-immunoprecipitation to explore the possibility of Cdca9 and Birc5b interacting in the early embryo. A mixture of *cdca9-flag* and *birc5b-gfp* mRNA was injected into a one-cell embryo and the injected-embryos were allowed to develop until sphere stage. Immunoprecipitation using an anti-Flag antibody was performed on the protein extracts from the injected embryos followed by Western blot analysis using an anti-GFP antibody. Under our conditions, Birc5b-GFP could be immunoprecipitated by anti-Flag antibodies, confirming that Cdca9 and Birc5b can interact in the early embryo (Supplementary Figure 3A).

### Aurora Kinase B activity is required for germ plasm RNP aggregation

To examine if the maternally expressed CPC requires its catalytic subunit, Aurora Kinase B, for germ plasm aggregation, we treated embryos with a known ATP competitive inhibitor for Aurora Kinase B, ZM-2. Past work in zebrafish has shown that the reduction of Aurora Kinase B function results in defects in furrow formation (43). Compared to DMSO-treated controls at 40 mpf, ZM-2-treated embryos showed a lower proportion of gpRNPs clusters (Figure 7; ZM-2-treated: 2-6 gpRNPs: 36.7%, ≥7 RNPs: 0.27%, N=1118, ROIs=30; DMSO-treated: 2-6 gpRNPs: 48.0%, ≥7 RNPs: 3.6%, N=1149, ROIs=30; Table 3) and a higher fraction of single-particle RNPs (ZM-treated: 63.3 %; DMSO-treated: 51.9 %). This result mirrors the reduction in germ plasm aggregation observed in *motley* and *cdca9* mutants, supporting that Aurora Kinase B activity in the context of the CPC plays a role in germ plasm aggregation.

**Figure 7:**
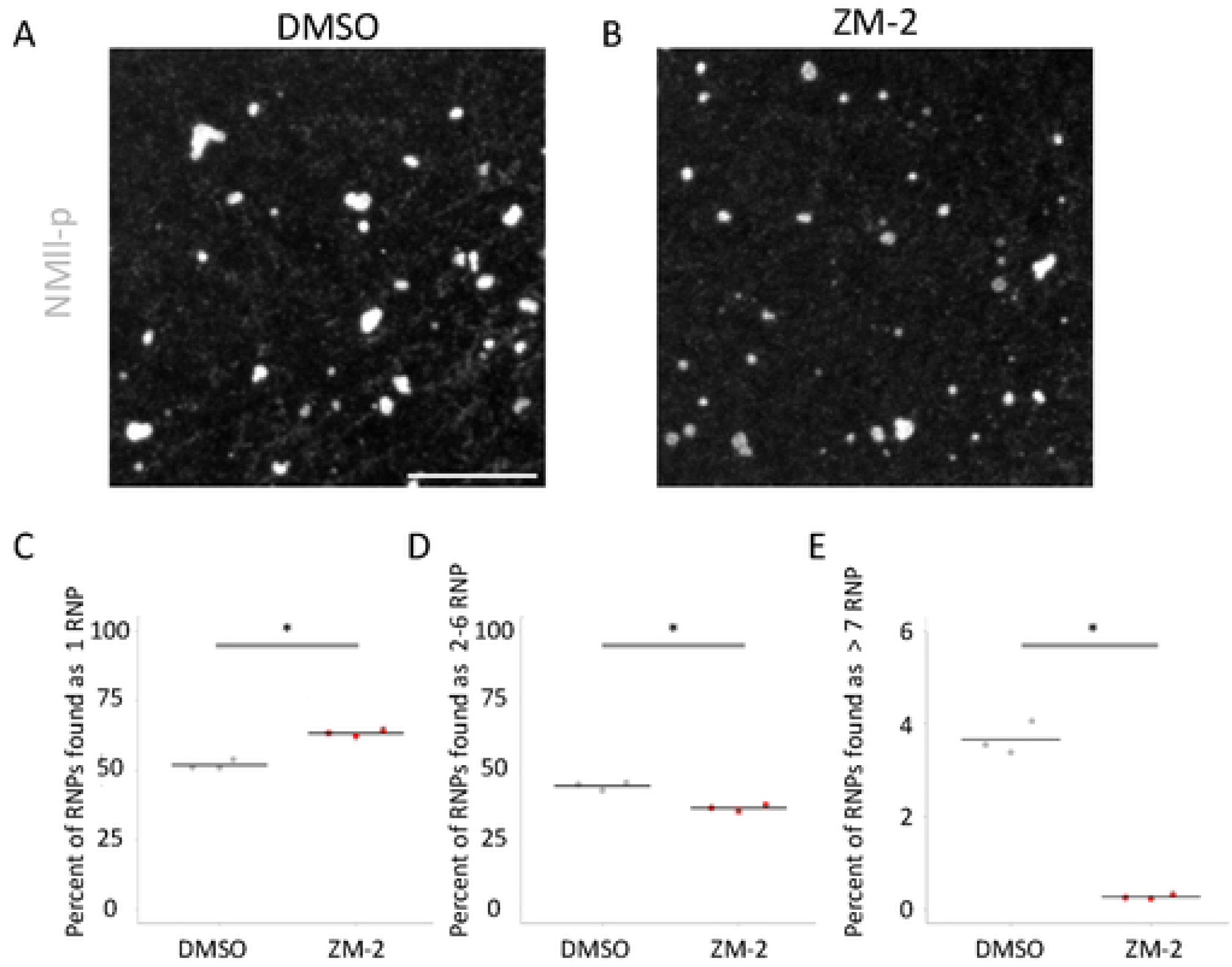
Aurora kinase B activity is necessary for the multimerization gpRNP. A-B) Images of gpRNPs at 40 mpf forming aggregates at the cortex prior to their recruitment at the furrow. A) DMSO-treated controls show gpRNP multimerization resulting in larger aggregates. B) ZM-2-treated embryos exhibit a reduction in the number of larger gpRNP aggregates compared DMSO-treated controls C-E) Quantification of aggregation according to size class, showing a significant increase in single RNPs and a decrease in the frequency of larger aggregates. Asterisks indicate statistical significance All micrographs are 2D maximum projections of confocal Z-stacks of animal views of blastodisc cortex, scale bars are 10 μm.

**Table 3:** Distribution of RNPs aggregate categories in ZM-2-treated embryos when compared to DMSO-treated controls.

| Genotype | 1 RNP | 2-6 RNP | >7 RNP |
| --- | --- | --- | --- |
| DMSO | 51.9 | 44.4% | 3.6% |
| ZN-2<br>P-value compared to DMSO | 63.3% | 36.4% | 0.27% |
|  | 0.0006 | 0.0017 | < 0.0001 |

### Birc5b interacts with germ plasm RNPs independent of the CPC

Our analysis shows the unique behavior of Birc5b as the single CPC component that accumulates with the growing gpRNP aggregates. We tested whether the interaction between Birc5b and gpRNPs is dependent on the CPC using Buc protein, which like NMII-P colocalizes with aggregating gpRNPs in wild-type (4), as an additional germ plasm marker (Figure 8A, ROIs=18). Interestingly, Birc5b and Buc continues to colocalize with Buc even in the *motley* mutant embryos, even though the truncated Birc5b protein, which contains only 79 amino acids of the N-terminal (with an extra 32 amino acids from an intron insertion for a total of 111 amino acids), lacks the coiled-coil domain required to form the triple helical bundle necessary for CPC stabilization (Figure 2, Supplementary Figure 3B-C) (Figure 8B ROIs=24, Supplementary Figure 2 A,B) (12,25). Similarly, Birc5b and Buc colocalize at the cortex in *cdca9* mutants (Figure 8C, ROIs= 24). Given the interdependence of CPC components for complex stabilization and targeting (25–27)(Fig. 6), these observations show that the interaction between gpRNPs and Birc5b relies solely on the N-terminus of the protein and is independent of CPC formation.

**Figure 8.**
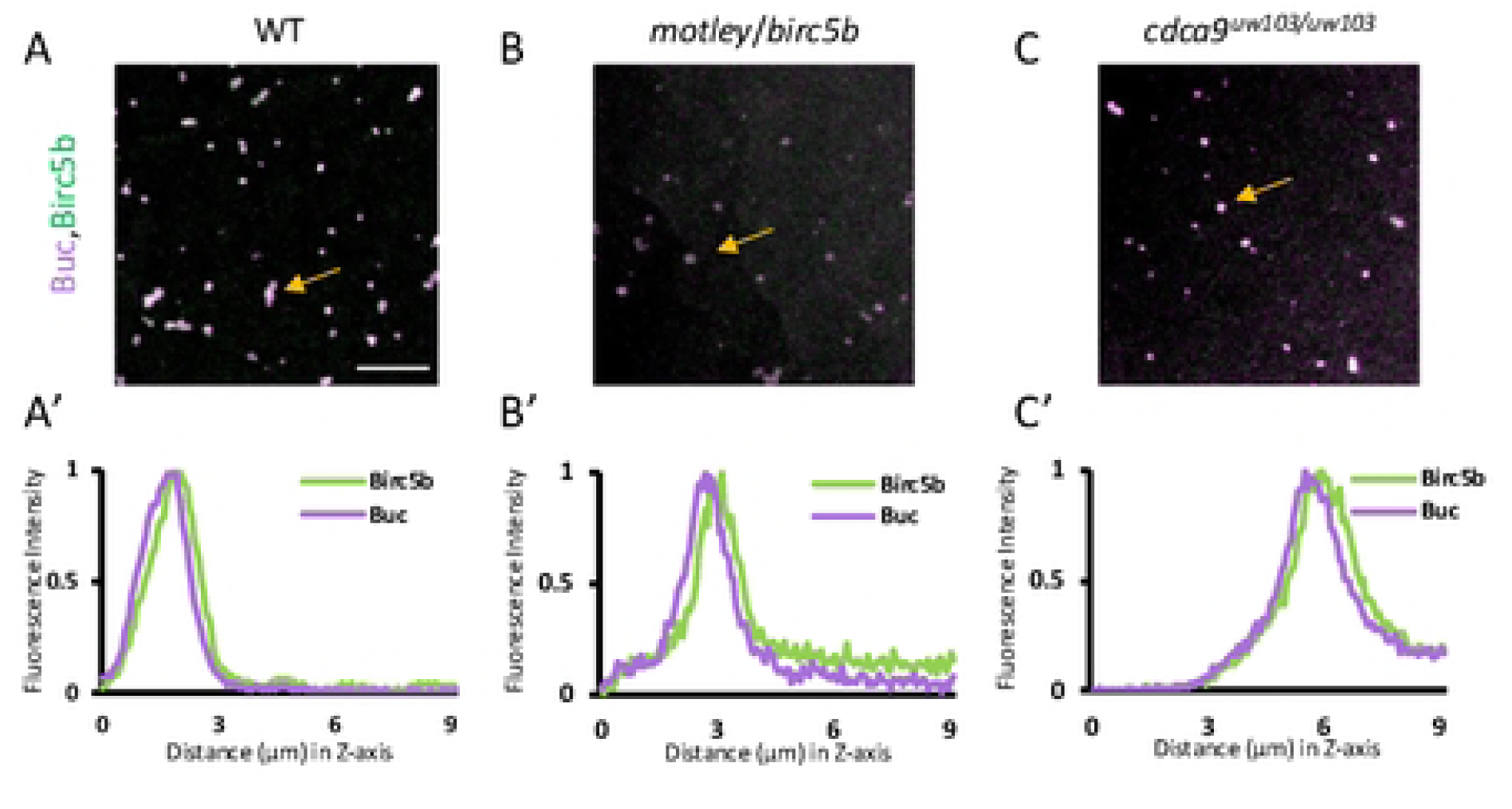
The interaction between Birc5b and gpRNPs is independent of the CPC. (A) Immunolabeling of Birc5b and Buc at the cortex of a wild type one-cell embryo shows accumulation of Birc5b in gpRNP aggregates. In *motley* and *cdca9* mutants (B,C), Birc5b and Buc continue to colocalize, showing that the interaction is independent of CPC formation. The gold arrow points to aggregates analyzed using relative fluorescence intensity in the Z-direction (A’-C’). All micrographs are 2D maximum projections of confocal Z-stacks, scale bars are 10 μm.

Collectively, our data suggest the presence of a maternally expressed CPC at the tips of astral microtubules that contains Birc5b, Cdca9, INCENP, and Aurora B kinase. This complex appears to behave similarly to the canonical CPC, with all four proteins essential for establishing a stable complex on the astral microtubules. At the same time and unexpectedly, Birc5b is able to interact with gpRNPs independently of CPC formation and accumulates within growing gpRNP aggregates.

### Maternally expressed CPC is necessary for germ plasm RNP attachment to microtubules

Previous research has revealed that the CPC can, directly and indirectly, interact with microtubules (44–49) and that the interaction between gpRNPs and microtubules is crucial for the aggregation of gpRNPs in the cortex of the embryo (12). To investigate whether the maternally expressed CPC is responsible for attaching gpRNP aggregates to microtubules, we examined the spatial relationship between gpRNPs and astral microtubules in the cortex of wild-type and the two CPC mutants.

Fixed embryos, from wild-type controls and the two CPC mutants, were imaged from the animal pole view to ascertain the spatial relationship between gpRNPs and astral microtubules in 3-D reconstructions (X-axis, along the plane of the cortex, and Z-axis, into the cortex) (Figure 9), by calculating the distance from the maximum intensity of the germ plasm RNP to the maximum intensity of the closest microtubule. This analysis showed that in wild-type embryos on average, cortical gpRNP aggregates are located approximately 0.68 μm from the nearest microtubule (Figure 9A,D,G, ROIs=16), a value similar to the previously reported estimated radius of single germ plasm RNPs (0.576 μm) (10) consistent with the direct apposition of gpRNP aggregates to astral microtubules. In contrast, gpRNP aggregates were not observed to be closely associated to astral microtubules in the cortex in the cortex of mutants for *cdca9* (Figure 9B,E,H) or *motley* (Figure 9C,F,I), with the nearest astral microtubule in the Z-axis in both mutants farther away from the gpRNP aggregates when compared to wild type (quantified in Figure 9J; *cdca9*: 1.87 μm ROIs=15, P-value < 0.0001; *motley*: 1.55 μm ROIs=15, P-value < 0.0001). Thus, maternally expressed CPC components are required for the attachment of gpRNPs to astral microtubules.

**Figure 9:**
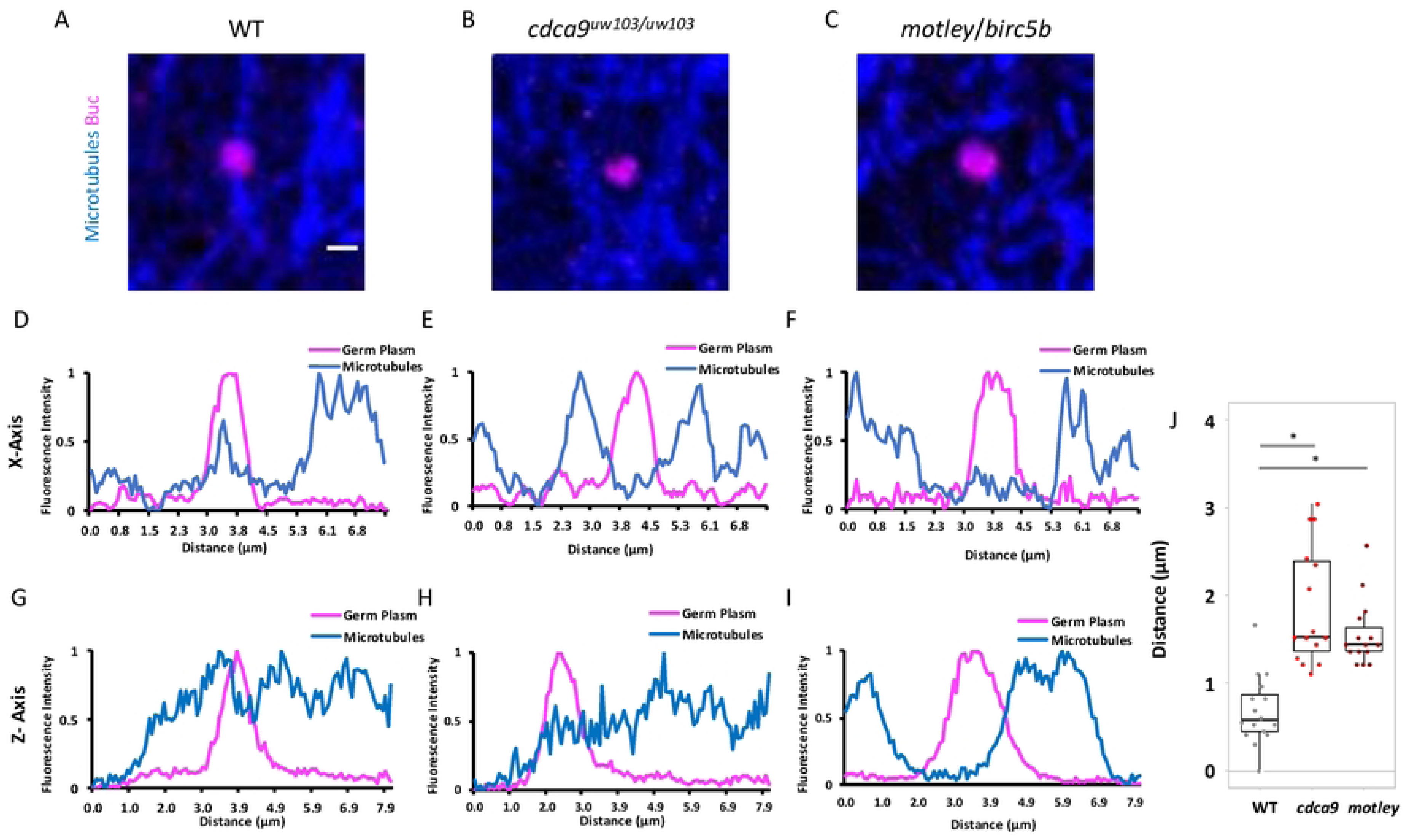
The maternally expressed CPC is necessary to attach gpRNPs to the microtubules. A-C) Micrographs from a 3D reconstruction visualizing the animal view of a 1-cell embryo labeled for microtubules and Buc to visualize germ plasm aggregates. In wild-type embryos (A), gpRNPs colocalize with microtubules. In *cdca9* (B) or *birc5b* (C) mutants, gpRNPs do not localize the microtubules. D-F) Relative fluorescence intensity in the X-Y direction (parallel to the surface of the cortex) across the germ plasm aggregates in (A-C). G-H) Relative fluorescence intensity in the Z-direction (going into the cortex) at the location of the germ plasm aggregate in (A-C). The germ plasm maximum is close to a microtubule maximum in wild type embryos (D,G) but not in *cdca9* (E,H) or *birc5b* (F,I) mutants. J) Dot plot of the distance between the closest maxima between microtubules and germ plasm in the Z-direction. Scale bars in (A-C) are 10 μm.

### Birc5b links gpRNP aggregates to furrow contractions during germ plasm distal compaction

In the early embryo, Birc5b is localized to the furrow and plays an essential role in the restructuring and bundling of microtubules during FMA formation at the developing furrow (12). To investigate if a maternally expressed CPC complex is present in the furrow with Birc5b, we examined the localization of other CPC members (Cdca9, Aurora Kinase B, and INCENP) during furrow ingression. A triple immunolabeling experiment, revealed that the bulk of Birc5b is localized to the furrow, while Cdca9 is confined to the edge of the furrow. Consistent with our findings prior to furrow formation, fluorescent intensity line scans across the forming furrow shows that microtubule maxima overlap with Cdca9 maxima, not Birc5b maxima (Figure 10A,B, ROIs=25). Similar to Cdca9, Aurora Kinase B and INCENP have intensity maxima at the edge of the furrow and not the furrow proper (Supplementary Figure 4)

**Figure 10:**
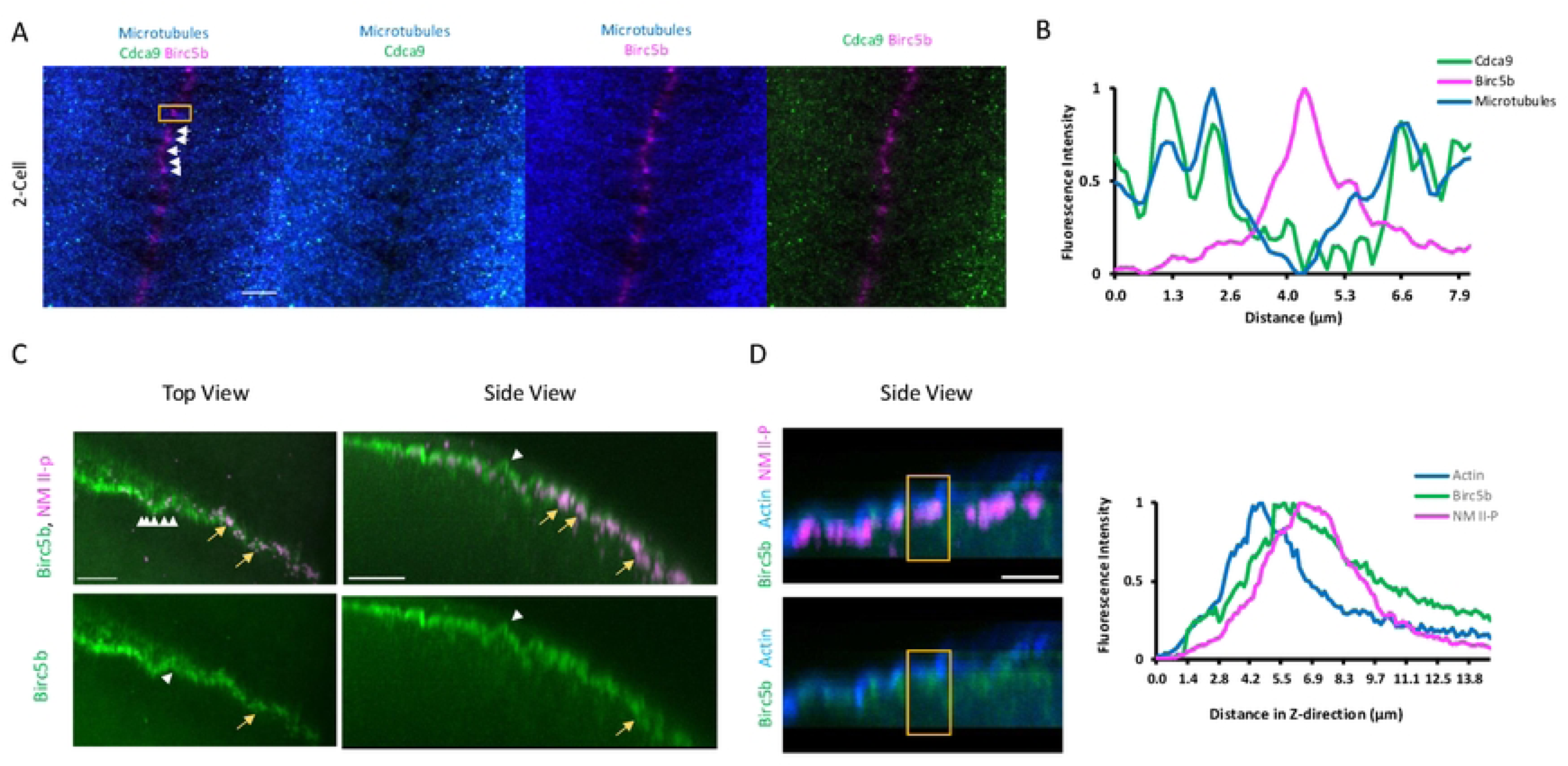
Birc5b is found with germplasm at the furrow. A) Animal views of an early 2-cell wild-type embryo at the region of the forming furrow labeled for Cdca9, Birc5b, and microtubules. Most Cdca9 localizes to microtubules in the regions at both flanks of the furrow whereas most Birc5b localizes to punctae along the furrow itself, at the center and associated with FMA microtubule bundles extending into the furrow from both sides of the furrow. Micrographs are 2D Maximum projections of confocal Z-stacks. B) A Line scan across the furrow (Gold box in (A)) shows that Cdca9 colocalizes with microtubules at both sides of the furrow while Birc5b is found at the furrow in a region devoid of microtubules. C) Top and side views of Birc5b and NMll-p, to label germ plasm aggregates, from a 3-D reconstruction of the furrow of a 2-cell stage embryo showing two patterns of Birc5b localization along furrow. A subset of Birc5b protein localizes with gpRNPs exhibiting the granular morphology characteristic of gpRNP aggregates (gold arrows). Another pattern of Birc5b localization can be discerned along the furrow in a ladder like pattern (white arrows) which corresponds to ladder-like bundling of microtubules (white arrows in A) as previously reported(12). The side view shows that the accumulation of Birc5b along the furrow exhibits a wave-like pattern D) A side view of Birc5b, NMll-p, and actin from a 3-D reconstruction of the furrow from a 2-cell embryo shows a similar wave like pattern for all components and the spatial relationship between all three proteins. A fluorescence intensity profile along the Z-direction shows the peak of Birc5b fluorescence intensity is observed between the actin and NMll-p, consistent with a role for Birc5b as a linker between actin and germ plasm. Scale bars are 10 μm.

To gain a greater understanding of the role of Birc5b in the furrow, we labeled two-cell embryos using anti-NMII-p antibodies to label gpRNPs and anti-Birc5b and created a 3D reconstruction of the furrow (Figure 10C, ROIs= 8). This analysis revealed that Birc5b exists in two distinct populations within the furrow, with a fraction of Birc5b at the inferred sites of microtubule bundling (as assessed by a ladder-like pattern, white arrow heads) in addition to a population of Birc5b associated with gpRNPs (gold arrows). Interestingly, the population Birc5b found at sites of microtubule bundling exhibits a pattern of repeated wave-like contractions along the furrow. A wave-like pattern has also observed when visualizing F-actin and it is thought to facilitate medial-to-distal furrow maturation including gpRNP aggregation towards the furrow’s distal ends (9,19). Additionally, similar to what is observed with F-actin (19), gpRNP-associated Birc5b is located within the grooves of the furrow contractions (gold arrows in Figure 10C).

To examine the spatial relationship between Birc5b, gpRNPs, and F-actin, we labeled these components in the furrow of an early two-cell embryo, performed a 3D reconstruction of the furrow and analyzed the fluorescence intensity in the Z-axis (going into the cortex). The results indicated that Birc5b is positioned between F-actin and gpRNPs at the developing furrow. Our data indicates that Birc5b is present at the interface between dynamic F-actin and germ plasm RNPs during furrow maturation as gpRNPs migrate to and become enriched at the furrow distal ends to form the germ plasm mass (Figure 10D, ROIs=5).

During furrow maturation, Cdca9 localizes to the furrow microtubule array and not to the distally compacted germ plasm masses (Figure 11 A,B, ROIs=9). In contrast, at the same stage Birc5b localizes with germ plasm as it undergoes compaction at the furrow distal ends (Figure 11 C,D, Supplementary Figure 5A,B) (NMII-p/Birc5b: 12/12; Buc/Birc5b: 14/14). Thus, our data indicate that the CPC is involved in gpRNP aggregation and furrow induction while localized to astral microtubule ends and the FMA, with the bulk of Birc5b remaining associated with aggregating gpRNPs as they undergo compaction at the distal ends of maturing furrows to form germ plasm masses.

**Figure 11:**
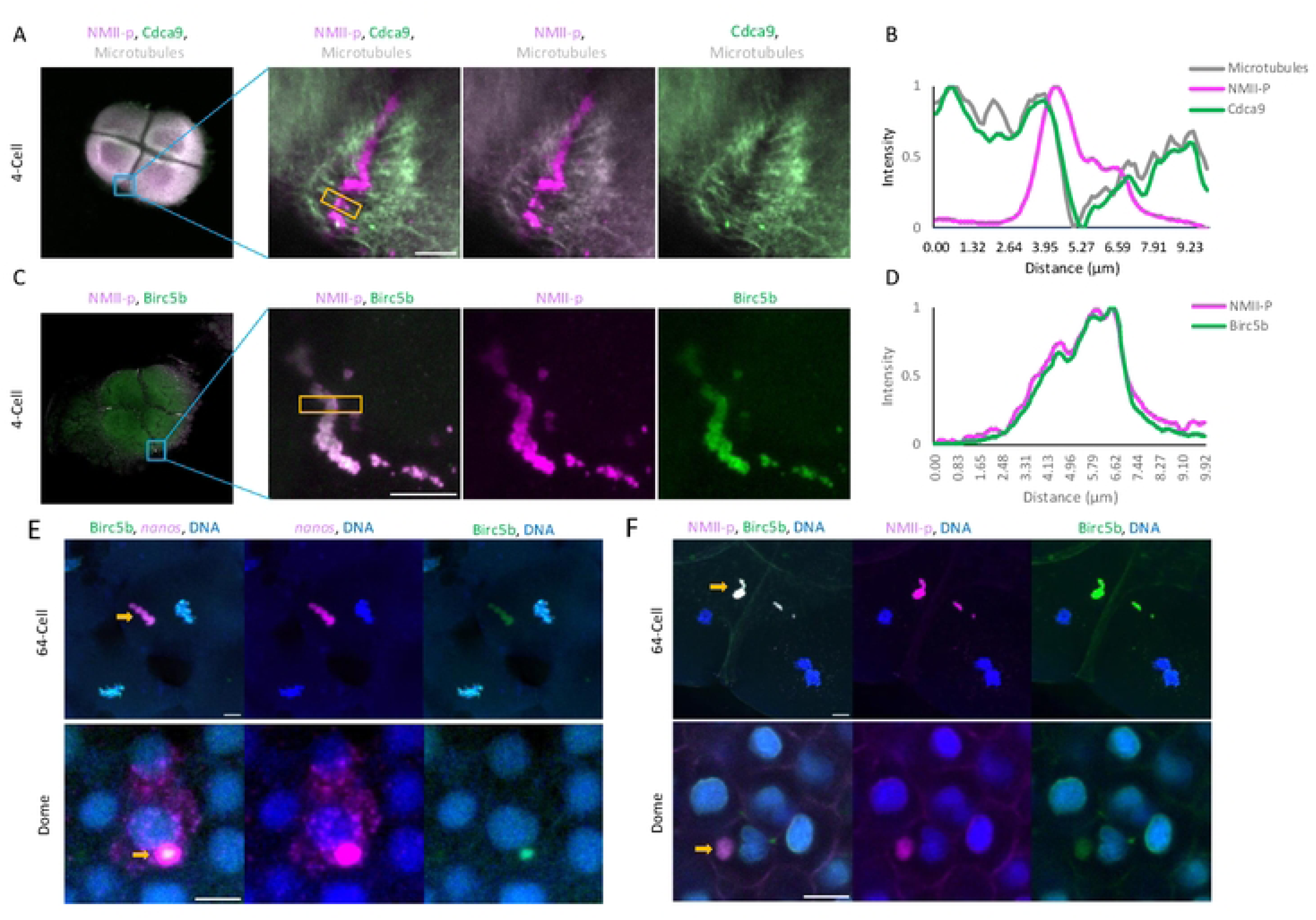
Identification of Birc5b within the germ plasm mass during early development. A) Micrographs visualizing the top view (blastodisc) of a 4-cell embryo labeled with microtubules, Cdca9, and NMll-p. The general localization pattern shows NMll-p localizes to the distal end of the furrows, while Cdca9 is localized to the FMA. B) A Line scan across the furrow (Gold box) shows that the majority of Cdca9 is colocalized with microtubules, not the germ plasm mass. C) Micrographs visualizing the top view (blastodisc) of a 4-cell embryo labeled with Birc5b and NMll-p. The general localization pattern shows NMll-p and Birc5b are colocalized to the germ plasm mass found in the distal end of the furrows. D) A Line scan across the furrow (Gold box) shows that the majority of Birc5b is colocalized with the germ plasm mass. E-F) Micrographs of germ plasm localization after it has become cellularized (64-cell stage) and during germ plasm RNA dispersal during the dome stage (∼4.3 hpf). E) germ plasm marker *nanos* RNA colocalizes with Birc5b protein at the 64-cell stage. At dome stage, *nanos* RNA disperses into the cytoplasm of the cell, while Birc5b remains in the germ plasm mass. F) Birc5b and NMll-p are colocalized together in the 64-cell stage, and during the dome stage, both proteins remain in the germ plasm mass. Micrographs are 2D Maximum projections of confocal Z-stacks, and scale bars are 10 μm.

### Birc5b remains associated with the germ plasm mass during early development

Previous studies showed that NMII-p, which colocalizes with gpRNPs during gpRNP recruitment and aggregation at the furrow for the first embryonic cycles (12), remains associated with germ plasm as it undergoes asymmetric segregation during the cleavage stages (50). Given the observed accumulation of Birc5b to aggregating gpRNPs, we tested whether Birc5b is also associated with germ plasm masses at later stages. We performed multiple time courses from 64-cell to dome, labeling both a germ plasm-associated RNA (*nanos*) and protein (NMII-p or Birc5b) (Figure 11E; Supplementary Figure 5 C) or labeling both proteins (NMII-p and Birc5b, Figure 11F). In all cases, we observe the colocalization of all three components, *nanos* RNA, NMII-p and Birc5b, within the germ plasm mass through cleavage stages and into early gastrulation, with no apparent spatial relationship between these components (Supplementary Figure 5 C; Supplementary Figure 6).

As development progresses and initiating during the early gastrula stages, individual gpRNPs undergo dispersal from the germ plasm mass into the cytoplasm, an event thought to help initiate the zygotic program for PGC specification (Eno et al., 2019; Knaut et al., 2000). Unexpectedly, at dome stage (about 4.3 hpf), when *nanos* RNPs undergo dispersal from the germ plasm mass into the cytoplasm, NMII-p and Birc5b continue to colocalize at an apparent remnant of the germ plasm mass (Figure 11 E,F, Supplementary Figure 5C).

Thus, both and NMII-p and Birc5b proteins appear distributed within the germ plasm mass during the cleavage stages, remaining associated to a germ plasm remnant even after gpRNPs undergo cytoplasmic dispersal.

### Differences between Borealin homologs

To gain insights into the function of Borealin homologs, we conducted a phylogenetic analysis of Borealin amino acid sequences from various vertebrate species (Supplementary Figure 7A). The identification of Cdca9 and Cdca8 across vertebrates revealed that Cdca9 is present in a subset of vertebrate genomes, specifically in several fish, amphibian, and reptile lineages (Supplementary Figure 7A), whereas it was not identified in invertebrate genomes. The distribution of Cdca9 in multiple distributed vertebrate linages suggest that it originated in an ancient gene duplication event in a vertebrate ancestor, prior to the later whole genome duplication at the base of the teleost lineage.

Phylogenetic analysis also showed that zebrafish Cdca8 is more closely related to mammalian Borealin protein than Cdca9. Comparing zebrafish Cdca8 to its human counterpart, we observed a 37% identity across the protein, while Cdca9 exhibited a lower identity of 21%. A protein domain analysis of Cdca8 identified the known N-terminal CPC interacting and C-terminal dimerization domains, with an intrinsically disordered region in between these motifs (Figure 1C, Supplementary Figure 7B)(51). Similar analysis of zebrafish Cdca9 shows the presence of the N-terminal CPC interacting domain and mid-region intrinsically disordered region (Supplementary Figure 7B) but does not identify a C-terminal dimerization domain (Figure 1C,).

Indeed, a comparison of the C-terminal regions of the zebrafish Cdca8 and Cdca9 proteins compared to human shows that Cdca9 exhibits less homology than zebrafish Cdca8 (17.9% and 43.4%, respectively) (Supplementary Figure 7C-E). Secondary structures of the three proteins predicts that Cdca9 contains an increased number of predicted ß-sheets when compared to both human and zebrafish Cdca8, although Cdca9 still contains conserved alpha-helix known to be involved in dimerization (52) (Supplementary Figure 7E).

## Discussion

Here, we show that *cdca9*, a maternally expressed homolog of borealin, is essential for early development in zebrafish embryos. Our characterization of *cdca9* function revealed that it not only has an expected role during meiosis and early embryonic mitosis, but also functions in the aggregation of cortical germ plasm gpRNPs in the early embryo (Figure 12). Our study indicates that a maternally-supplied CPC that includes Birc5b and Cdca9, products of gene duplicates for the known canonical CPC components *birc5a/survivin* and *cdca8/borealin*, acts as a linker between microtubules and gpRNPs in the cortex to promote aggregation prior to and during furrow formation. Unexpectedly, as gpRNPs aggregate even prior to furrow recruitment, Birc5b, but not other CPC components, remains associated with gpRNP aggregates. The association of Birc5b with gpRNPs continues during distal compaction and as germ plasm masses are asymmetrically inherited during cell division until gpRNP dispersal at gastrulation.

**Figure 12:**
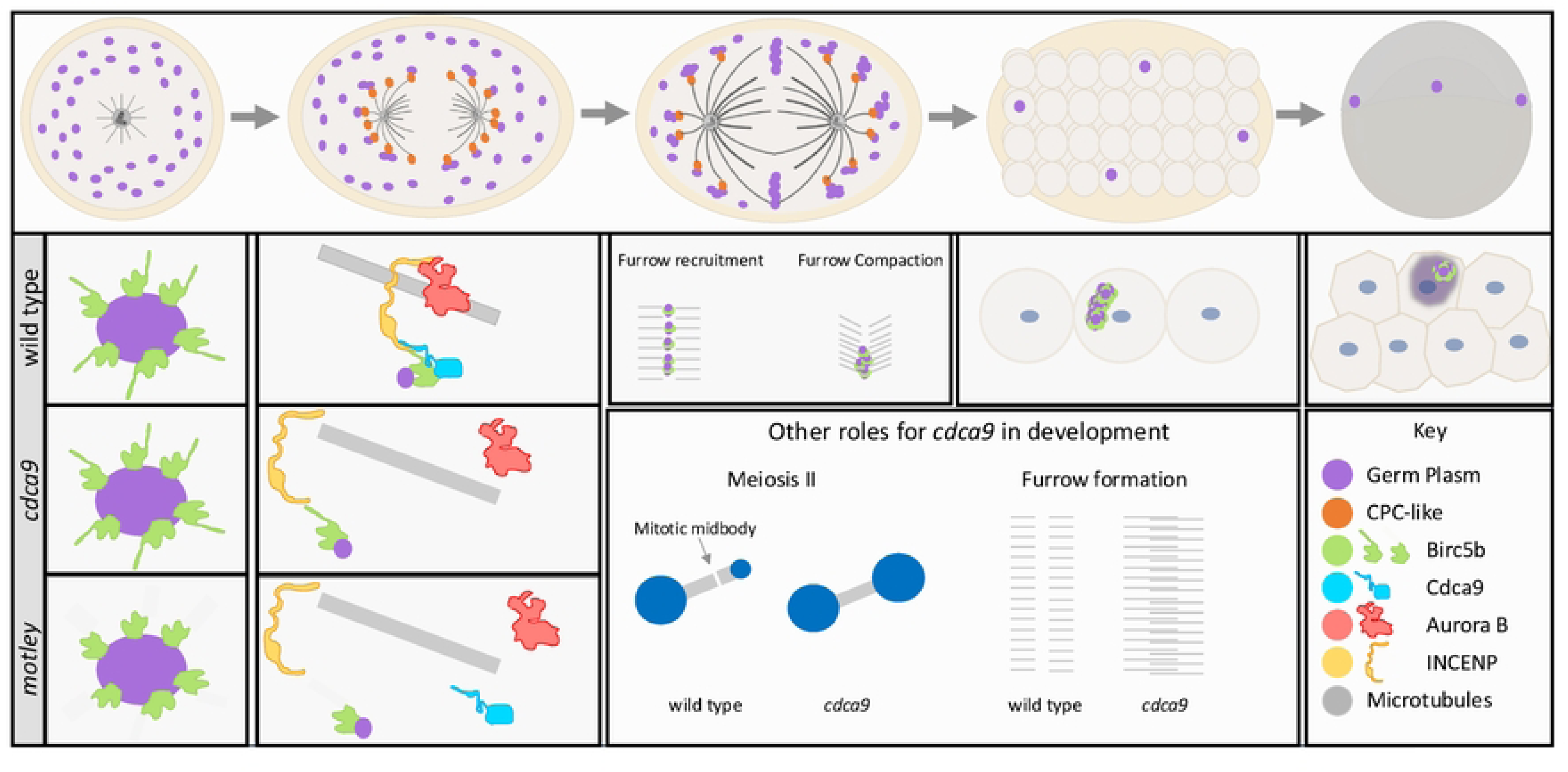
Model highlighting multiple roles for Cdca9 and Birc5b in germ plasm organization. In wild-type embryos, Birc5b colocalizes with gpRNPs in the cortex prior to the first mitosis; this interaction is not dependent on a maternally expressed CPC. As the astral microtubules grow out, maternally expressed CPC containing Birc5b and Cdca9 is required to attach the aggregating gpRNPs to the ends of the astral microtubule. When either Birc5b or Cdca9 is missing, the gpRNPs cannot attach to the microtubules. During the first embryonic cell division, gpRNP/ Birc5b aggregates are recruited bilaterally into the forming furrow, and the association between gpRNP and Birc5b continues through distal compaction and after the germ plasm mass has become cellularized. During early cleavage stages, the germ plasm mass that contains Birc5b is inherited asymmetrically. In the last blastula/early gastrula embryo, germ plasm masses undergo cytoplasmic dispersal of germ plasm mRNAs but not Birc5b, which remains in a germ plasm remnant. Cdca9 is also required during meiosis for midbody formation during polar body extrusion and during early embryonic mitoses to establish the furrow, the latter including severing function of astral microtubules as they reach the furrow and their reorganization into the FMA.

### Role of maternally inherited CPC in cell division during meiosis and early cell division

Our studies indicate that Cdca9 and Birc5b can create a specialized maternally inherited CPC necessary for cellular division during early development. In other species, the CPC has been found to promote spindle assembly in both mitotic and meiotic cells (33,39–41,53,54). When CPC factors have been depleted from cells, a reduction in spindle length is typically observed (12,53,55). In the case of the *motley* and *cdca9* mutants, we detect shortened spindles in both mitotic and meiotic cells, respectively (12) (These studies). In human oocytes, Cdca8 has been found to be localized to the meiotic spindle, with higher levels during meiosis II, and its depletion hampers the rate of polar body exclusion although it does not block oocyte development (41). This is consistent with what we observe in both *motley* and *cdca9* mutant eggs, which exhibit defects in DNA segregation and/or in the condensation of the polar body nucleus indicative of defective or slower polar body extrusion during meiosis II, yet the eggs are still able to be fertilized (12).

During mitosis, the CPC has multiple roles during cellular division, including regulating kinetochore attachment errors, furrow ingression, and abscission in the cell (24,56). Consistent with these roles, both *motley* and *cdca9* mutant embryos fail to create furrows but are still able to undergo multiple rounds of DNA replication and segregation, albeit exhibiting aberrant DNA morphology (12) (These studies). During early development, the zebrafish embryo does not have cell cycle checkpoints (57), which may allow DNA replication and segregation to occur with improper attachments, leading to uneven DNA segregation. Additionally, cytokinesis defects in *motley* and *cdca9* mutants may also contribute to uneven DNA segregation, given the lack of membranes to insulate chromosome sets from the influence of adjacent bipolar spindles. The phenotypes observed in the *motley* and *cdca9* mutants suggest that a specialized maternally-expressed CPC is required for cell division in the early zebrafish embryo, in a similar manner as the canonical CPC.

### A maternal CPC has a role in germ plasm RNP aggregation in the cortex

Our studies indicate a new role for a maternally-expressed CPC in the early embryo, in the aggregation of gpRNPs at the cortex prior to and during furrow formation for the first several cell cycles. We find that the removal of either Cdca9 or Birc5b function from the early embryo results in significantly impaired gpRNP aggregation when compared to wild type (12) (These studies). Additionally, embryos treated with an Aurora kinase B inhibitor also showed a decrease in germ plasm aggregation, although further studies will determine the precise role of Aurora kinase B activity in this process.

Previous research involving functional and subcellular localization data suggest that gpRNP aggregation in the cortex of the early embryo depends on the growth of astral microtubules whose plus ends are bound to gpRNPs, contributing to the radially outward movement of the latter towards the furrow (12,14). Moreover, Birc5b has been proposed to act as a necessary linker between astral microtubule ends and gpRNP aggregates (12). Similar to Birc5b, we find that absence of Cdca9 function results in gpRNP aggregation defects and the lack of association of germ plasm RNP aggregates to microtubules.

Immunolabeling of CPC members is consistent with the presence of a complex on the tips of the astral microtubules that contain Birc5b, Cdca9, INCENP, and Aurora B kinase, with colocalization dependent on Birc5b and Cdca9 function. This agrees with the known structure of the conventional CPC, where Borealin, Survivin, and INCENP directly interact with each other, creating a triple helix that cannot be stabilized without inter-dependent interactions from other CPC members (25,33,35,36). We also find that overexpressed Cdca9 and Birc5b can interact in embryonic extracts. Altogether, our data indicates that Cdca9 and Birc5b work together within the context of a novel maternally inherited CPC to mediate gpRNP aggregation, by linking the action of growing astral microtubules to gpRNP multimerization.

### Birc5b interaction with Germ Plasm RNP is independent of the CPC

In spite of the known interdependence of CPC stabilization on its four members, we unexpectedly find the interaction between Birc5b and germ plasm aggregates is not dependent on CPC formation. In particular, a Birc5b product corresponding to the *motley* mutant variant, containing a truncated N-terminal BIR domain is still able to associate with gpRNPs aggregates (12)(Theses studies) in spite of lacking a C-terminal coiled-coil domain necessary for interactions with INCENP and Borealin (25) (These studies). Thus, the interaction between Birc5b and gpRNP depends on the N-terminal-most region of Birc5b, does not require its full BIR domain, and is independent of CPC formation.

Also unexpectedly, once gpRNP aggregates reach the furrow, only Birc5b, but not other CPC members, maintains its association with gpRNPs as they continue to undergo segregation and become compacted at the distal end of maturing furrows. Moreover, during furrow maturation gpRNPs are observed between F-actin indentations, when repeated contractions in F-actin along the furrow facilitates furrow maturation and the distally-directed compaction of germ plasm masses (9,19). We find that during this process a gpRNP-colocalizing Birc5b population is also associated with furrow F-actin indentations. Moreover, Birc5b at the furrow localizes between furrow actin and gpRNP aggregates, consistent with a potential role for Birc5b in anchoring gpRNPs to the F-actin cytoskeleton at the furrow. Such anchors would allow transducing the movement of distally-directed F-actin contraction waves into the distally-directed movement of aggregating gpRNPs, mediating gpRNP accumulation and coalescence as a compacted germ plasm mass at furrows distal ends. Previous studies have shown that Birc5b is necessary to organize F-actin in the cortex of the early embryo, which is associated with gpRNPs (12) and we can not rule out a role for Birc5b in mediating F-actin dynamics at the furrow.

### Birc5b is localized to a biomolecular condensate

Phase-separated condensates are formed when different biomolecules, such as proteins or nucleic acids, separate from the cellular environment as a distinct fluid phase, creating membrane-less organelles. Recently, it has been proposed that the germ plasm mass in zebrafish is a biomolecular condensate. This is supported by findings that treatment of zebrafish embryos with hexanediol results in germ plasm mass fragmentation, consistent with germ plasm possessing hydrogel properties, and that Buc protein homologs exhibiting fluid phase properties in vitro (6,58,59).

Our studies indicate that, in addition to NMII-p, Birc5b integrates into the germ plasm biocondensate independent of other CPC components (These studies)(50). Although biomolecular condensates can have an internal organization related to their composition and cellular activities (60), neither NMII-p nor Birc5b exhibits a discernable spatial distribution within the germ plasm aggregate. After the germ plasm RNA undergoes dispersal during the early gastrula stage, however, NMII-p and Birc5b do not disperse and instead remain as an apparent germ plasm remnant. Further studies will be needed to elucidate the role of Birc5b and NMII-p in the segregation and dynamic restructuring of the germ plasm aggregate during early zebrafish development. Such potential roles include contributing to the structure and/or subcellular segregation of the germ plasm biocondensate in the early embryo.

### Subfunctionalization of a maternal-specific CPC

We find that, in zebrafish, both Survivin and Borealin genes exhibit duplicated copies, with one set of gene duplicates (*birc5a* and *cdca8*, respectively) expressed throughout development and another set (*birc5b* and *cdca9*) exhibiting maternal-specific expression. Moreover, ubiquitously expressed genes *birc5a* and *cdca8* show greater sequence similarity to corresponding genes in species without a gene duplication, while *birc5b* and *cdca9* exhibit greater divergence (12) (These studies). Thus, zebrafish *birc5a* and *cdca8* likely carry out general functions required across the organism’s lifespan, whereas *birc5b* and *cdca9* appear to be duplicate copies that have subfunctionalized to have a maternal-specific function. On the other hand, zebrafish INCENP and Aurora Kinase genes, also components of the CPC, occur as single copies. Thus, subfunctionalization of a multiprotein complex, the CPC, to carry out functions specifically in the very early embryo appears to be achieved through the duplication and divergence of some, but not all, components of the complex.

In conclusion, our analysis highlights the essential role of maternally expressed homologs of CPC members *borealin* (*cdca9*) and *survivin* (*birc5b*) in early development, within the context of a maternal-specific CPC. As expected from the known function of the conventional CPC, these maternal CPC components are involved in the regulation of cell division at the egg to embryo transition. Unexpectedly, they also play a crucial role in germ plasm RNP aggregation in the early embryo, acting as a linker between microtubules and gpRNPs at the cellular cortex. We also identified a novel association of Birc5b, but not other CPC components, with germ plasm RNPs as the latter become compacted to generate germ plasm masses during furrow formation and undergo segregation in the cleavage stages, until RNP cytoplasmic dispersal during gastrulation. Our studies reveal a key role for the CPC in the inheritance and segregation of germ plasm determinants.

## Material and Methods

### Animal husbandry

Stocks of wild-type (AB), *motley* (*mot^p1aiue^)*, and *cdca9* mutant fish lines were raised and maintained under standard conditions at 28.5°C (61). Embryos were collected by natural matings and allowed to develop in E3 medium, with staging according to developmental time and morphological standards (62). All zebrafish housing and experiments were approved by the University of Wisconsin-Madison (assurance number A3369-01).

### Genetic identification of homozygous mutant fish

Genomic DNA were extracted by either placing a clipped piece of tail fin or a single embryo in 100 ul of 50 mM NaOH and boiled at 95°C for 20 minutes. After the samples were cooled down to 4°C, 10 ul of Tris-HCl pH 8.0 was added to neutralize the lysate.

Homozygous *motley* (*mot^p1aiue^)* fish were identified using the locked nucleic acid (LNA) endpoint genotyping method with the HEX fluorescent probe identifying the wild-type allele and the FAM fluorescent probe identifying the mutant allele. LNA endpoint reactions were performed using PrimeTime Gene Expression Master Mix (IDT) at a final volume of 10 ul with 250 nM of each probe, 1000 nM of each primer and 10-25 ng of DNA. Fish homozygous for CRISPR-Cas9 generated *cdca9* mutant alleles were detected via PCR, employing a 100 bp primer fragment over the CRISPR-Cas9 target site, and the PCR product was resolved on a 2.5% agarose gel. Detailed primer and probe sequences for each mutant can be found in Table 1. Genotypes were additionally verified by observing the expected maternal-effect phenotype in the offspring.

### CRISPR-Cas9 mutagenesis of *cdca9*

The CRISPR design web tool CHOPCHOP was used to identify a guide RNA site within the first active domain of the *cdca9* as identified by Ensembl (63–65). A Guide RNA template was produced using an annealing/fill-in method (66). Briefly, two oligos, first a gene-specific oligo containing a T7 promoter, the gene specific target sequence and a complementary region to the constant oligo, and second, a constant oligo encoding the reverse complement of the tracrRNA tail, were annealed together and ssDNA overhangs were filled in using T4 DNA polymerase (NEB). The resulting sgRNA template was purified using DNA Clean and Concentrator Kit (Zymo Research). Guide RNA was created using the MEGAshortscript T7 Transcription Kit (Invitrogen) and purified using an ethanol/ammonium acetate protocol. The concentration of the guide RNA was measured using a NanoDrop spectrophotometer, and integrity was checked by gel electrophoresis. The guide RNA was stored as single-use aliquots at −80°C.

To generate a targeted mutation, a mixture of sgRNA (final concentration: 200 pg/nl) and Cas9 protein (PNABio, final concentration: 400 pg/nl) was injected into the blastodisc of one-cell wild-type embryos. At 24 hpf, a subset of injected embryos was collected, and DNA was extracted as described above. To confirm CRISPR-Cas9 activity in somatic tissues, PCR was carried out using the genotyping primers described above and the resulting products were resolved on a 2.5% agarose gel to visualize DNA changes, visible as a smear in the gel (67,68) (Table 1).

### Transcript expression

Total RNA was extracted from pools of five wild-type embryos using Trizol (Invitrogen) according to manufacturer instructions. cDNA was synthesized using the Super Script III reverse transcriptase kit (Invitrogen) with a random hexamer primer. RT-PCR was performed using primers found in Table 1 and resolved on a 2.5 % agarose gel with β-actin acting as a control.

### Cloning of Cdca9

Total RNA extraction and cDNA synthesis were performed as described above, except that an oligo-dt primer was used to generate full length cDNA transcripts instead of a random hexamer primer. Full length coding transcripts were amplified from cDNA library using primers found in Table 1. The PCR products were ligated into pCR 2.1 vector (Invitrogen) and verified by sequencing. The full-length coding transcripts were subcloned into pCS2Flag (Addgene plasmid # 16331) expression vector, a gift from Peter Klein. *cdca9* transcripts were inserted into the pCS2 expression vector at the Cla1 site using the In-Fusion Snap Assembly mix (TakaRA) and site-specific primers (Table 1). The final constructs were confirmed by Sanger sequencing. Sense mRNA for the immunoprecipitation experiments was synthesized from the SP6 promoters using the mMessage mMachine kit (Invitrogen) and Not1 linearized plasmid DNA.

### Immunoprecipitation

*cdca9-flag*, or *birc5b-gfp*b (12) were injected into one-cell embryos at a final concentration of 200 pg/nl of each mRNA. The injected embryos were allowed to develop for 4 hours, then 100 injected embryos were transferred into a non-stick microfuge tube. Then embryos were deyolked using a deyolking buffer consisting of 55 mM NaCl, 1.75 nM KCl, and 1.25 mM NaHCO_3_ and the embryo pellet was snapped frozen until needed.

On the day of the immunoprecipitation experiment, the frozen embryo pellet was homogenized in lysis buffer containing 10 mM Tris-HC1 pH 8, 150 mM NaCl, 1 mM EDTA pH 8, 1% Triton X-100, and 1x Halt Protease Inhibitor Cocktail (Thermo Fisher Scientific-78430), the embryonic extract was spun at maximum speed in a microcentrifuge (>20,000 x g) at 4^0^C for 10 minutes, and the supernatant was transferred to a new non-stick microfuge tube. Protein concentration was quantified using a spectrophotometer (Eppendorf BioPhotometer). In each experiment, 2000 ng of input was incubated with ANTI-FLAG M2 Affinity Gel (Sigma A2220) overnight at 4°C. The next day, the beads were washed 3 times in Tris-buffered saline (TBS). To elute the sample, 20 μl of 2X Laemmli sample buffer (Bio-Rad −161073) were added to the tube and the mixture was heated for 10 minutes at 50°C. After heating, samples were centrifuged for 30 seconds to pellet any undissolved agarose and the supernatant was transferred to a fresh non-stick microfuge tube.

Samples were loaded on a precast 4-15% TGX gel (Bio-Rad) and blotted to a Immobilon-FL PVDF membrane (Millipore) at 1 hour at 100V. Membranes where blocked using Intercept Blocking buffer and incubated with 1:2000 mouse anti-Flag M2 antibody (Sigma-Aldrich F1804), or 1:400 Rabbit anti-Birc5b (12). Membranes were developed using IRdye800 CW goat anti-Mouse Secondary Antibody (1:15000) or IRDye-680RD goat anti-Rabbit secondary antibody (1:15000). The processed blot was visualized using the Odyssey Imager system.

### ZM2 drug exposure

ZM2 (ZM44739, Tocris Bioscience) was dissolved in DMSO at a concentration of 20 mM. Dechorionated wild-type embryos were exposed to 400 µM ZM2 diluted in embryonic medium starting at 15 minutes after fertilization until fixation at 40 minutes after fertilization.

### Immunolabeling

Embryos from all fish lines were collected and dechorionated with a Pronase solution (Roche-10 165 921 001) and placed into E3 embryonic medium until they were fixed at the appropriate time point in cytoskeleton-preserving fixative (4% paraformaldehyde, 0.25% glutaraldehyde, 5 mM EGTA, 0.2% Triton X-100 in phosphate buffered saline (PBS)) overnight at 4°C. Fixed embryos were permeabilized with methanol and stored at −20°C until use. Embryos were rehydrated through a methanol:PBS series, treated with 0.05mg/ml sodium borohydride solution for 30 minutes at room temperature to inactivate the glutaraldehyde, and subsequently washed with PBS-Triton. Embryos were heated at 68°C for three hours, blocked for one hour in Blocking Reagent (Roche 1109617600), and incubated in primary antibody overnight at 4°C. The next day, the embryos were washed with PBS-T and incubated in secondary antibodies overnight at 4°C. To detect DNA, embryos were labeled with 0.5 μg/ml solution of 4′,6-diamidino-2-phenylindole-dihydrochloride (DAPI) for 10 min followed by three washes in PBS. After labeling embryos were deyolked and mounted flat with the animal pole upwards in 75% glycerol.

Antibodies against Cdca9 were derived from the C-terminus sequence (Abmart, X-P86347-C), used as unpurified samples and validated by lack of immunofluorescence labeling in *cdca9* mutant whole mounts. Two antibodies were used interchangeably to label germ plasm, Rabbit anti-Bucky Ball and Rabbit anti-phospho-Myosin Light Chain 2 (12,69). Primary antibodies used for immunolabeling of fixed embryos were as follows: rabbit anti-β-catenin (1:1000, C2206, Sigma-Aldrich), mouse anti-α-Tubulin (1:2500, T5168, Sigma-Aldrich), rat anti-α-Tubulin (1:1000, ab6161, Abcam), Rabbit anti-phospho-Myosin Light Chain 2 (1∶50, Cell Signaling Technology 3671L), Rabbit anti-AurB (1∶100)(43), Rabbit anti-Survivin Alexa 488 (1∶25, Cell Signaling Technology 2810), Rabbit anti-Survivin Alexa 647 (1∶25, Cell Signaling Technology 2866), Rabbit anti-INCENP (1∶25, Invitrogen PA5-17200), mouse anti-Cdca9 (1:250), and Rabbit anti-Bucky Ball (1:250,(69)).

To detect actin and other proteins in colocalization experiments, dechorionated embryos were fixed in 4% paraformaldehyde, 0.25% glutaraldehyde, 5 mM EGTA, 0.2% Triton X-100 and 0.2 U/ml rhodamine-conjugated phalloidin for 3-5 h at room temperature and then overnight at 4°C. After the overnight incubation, the fixed embryos were then treated with sodium borohydride solution and proteins detected through immunofluorescence as above.

### Combined immunofluorescence and *in situ* hybridization

For combined immunofluorescence and *in situ* hybridization, embryos were fixed in cytoskeleton-preserving fixative as described above. Fluorescent *in situ* hybridization was performed with a Cy3-labeled antisense *nanos* probe (Köprunner et al., 2001). DIG haptens were detected using the TSA Plus CY3 system (PerkinElmer) following a modified “Triple fluorescent in situ” protocol (71). After the *in situ* hybridization the embryos were washed in PBS-Tween and blocked in Blocking Reagent (Roche 1109617600). The embryos were then incubated with primary and secondary antibodies, labelled with DAPI to detect DNA and deyolked and mounted as described above.

### Image acquisition and processing

Embryos were imaged using a Zeiss LSM510, Zeiss LSM780 or Zeiss LSM980 confocal microscope. Microscope optics used includes EC Plan-Neofluar 10X/0.30 M27, C-Apochromat 40x/1.10 W corr M27, C-Apochromat 63x/1.2 W corr M27, Plan-Apochromat 63x/1.40 Oil DIC M27, and alpha Pan-APO 100x/1.46 oil DIC M27. Images were processed using standard image processing in ImageJ (72) including maximum 2D Z-projection of multiple slices from 3D stacks, with adjustments to brightness and contrast applied across the entire image and similarly across samples. To determine colocalization, the plot profile and plot z-axis profile tools were used. In each experiment, the fluorescent intensity value in the same area was acquired for each channel and normalized so that the maximum fluorescent intensity value of each channel was equal to one and the minimum intensity value was equal to zero. The Coloc2 plugin was used to estimate the colocalization between CPC components using Pearson’s correlation coefficient. As a colocalization control, the Pearson’s correlation coefficient value corresponding to random colocalization was determined by rotating a single channel of the image and comparing it to the untransformed channel (73). All image data analysis was performed on a minimum of 3 embryos per condition.

## Acknowledgments

We thank past and current members of the Pelegri lab for their contributions to our research, particularly our animal husbandry staff for the care of the aquatic facility. We are also grateful for the advice and protocol provided by Dr. Edlyn Wu for immunoprecipitation. Microscopy was performed at the Newcomb Imaging Center, Department of Botany at the University of Wisconsin-Madison.

---

**Supplementary Figure 1:**
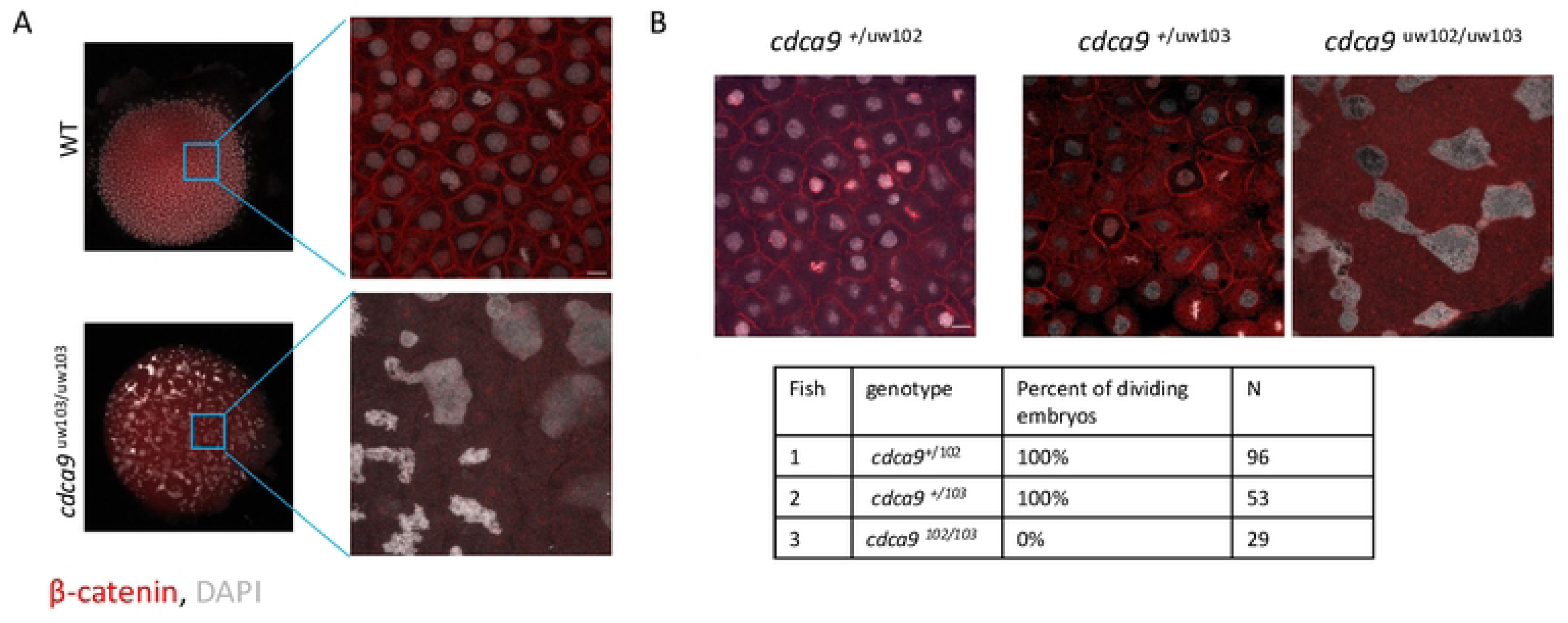
Non-complementation of the two CRISPR-Cas9 *cdca9* alleles. A) Micrographs visualizing cell adhesion junction component ß-catenin and DNA (DAPI, gray) show discrete cell membranes with a nucleus in all cells in the wild type. In *cdca9* maternal mutants, there is no ß-catenin accumulation, reflecting the lack of membrane formation, and DNA clusters appear as irregular, larger aggregates. B) Embryos from cdca9 heterozygous females (*cdca9^102^*^/+^ or *cdca9^103^*^/+^) have mature cell membranes marked by ß-catenin with a nucleus in all cells. In contrast, embryos from transheterozygous *cdca9*^102/103^ females show no ß-catenin accumulation, and DNA appears in irregular, larger aggregates. All micrographs are 2D maximum projections of confocal Z-stacks, and scale bars are 10 μm.

**Supplementary Figure 2:**
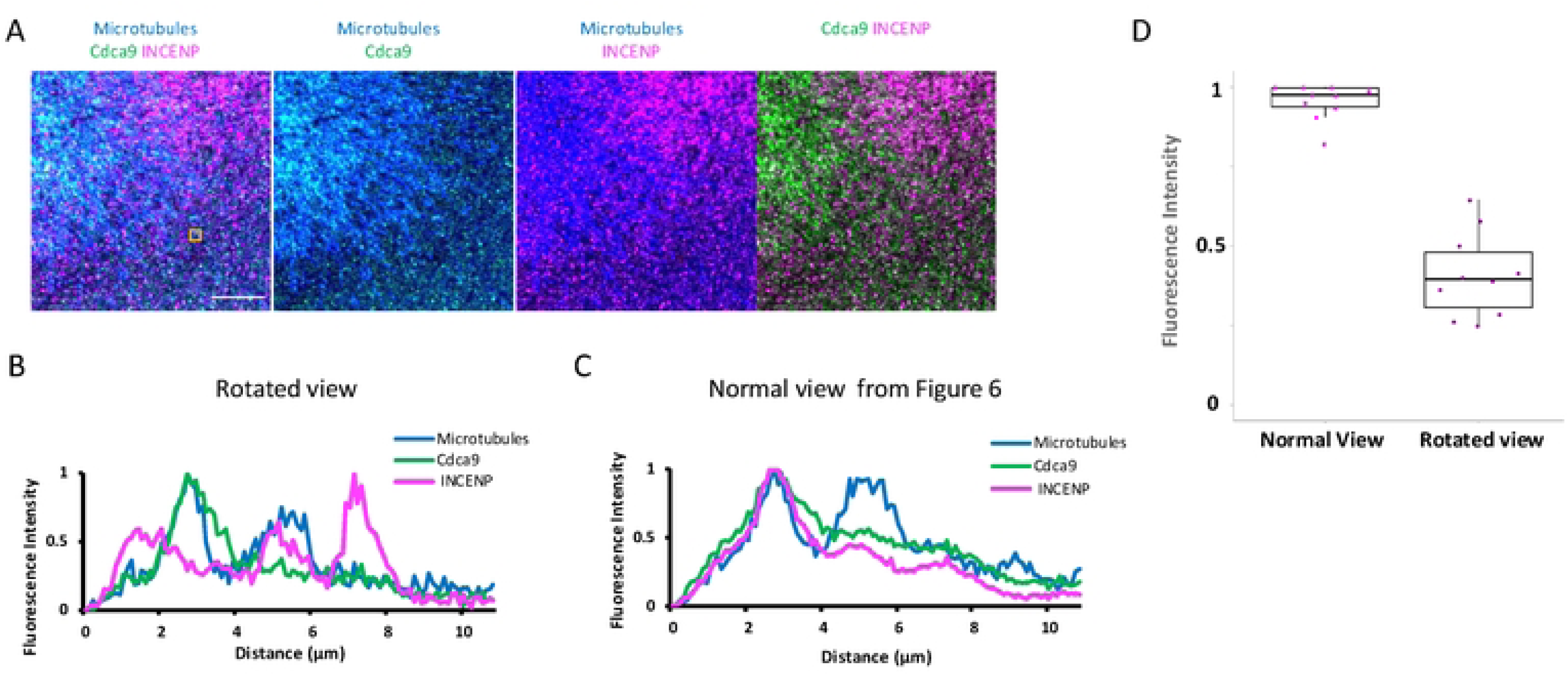
Rotated control for Microtubules Cdca9 and INCENP micrograph. A) The micrograph from Fig. 6 visualizing the astral microtubules in a one-cell embryo showing the location of Cdca9 and INCENP with the INCENP channel being rotated 90 degrees as a control for randomization. B) The intensity profile of a Cdca9 puncta, as marked by the gold box, shows relative fluorescence intensity in the Z-direction, showing that Cdca9 and INCENP do not colocalize in the rotated view micrograph. C) Intensity profile of the Cdca9 INCENP puncta from Fig 6 as a comparison. D) Dot plot showing the relative fluorescence intensity of INCENP at the maximum relative fluorescence intensity of the Cdca9 punctae in both the regular and rotated view, with significantly less relative fluorescence intensity of INCENP in rotated view (regular view: mean-0.955 ROIs:10, rotated view: mean-0.409 ROIs:10, P-value > 0.0001). Micrographs are 2D maximum projections of confocal Z-stacks, and scale bar is10 μm.

**Supplementary Figure 3:**
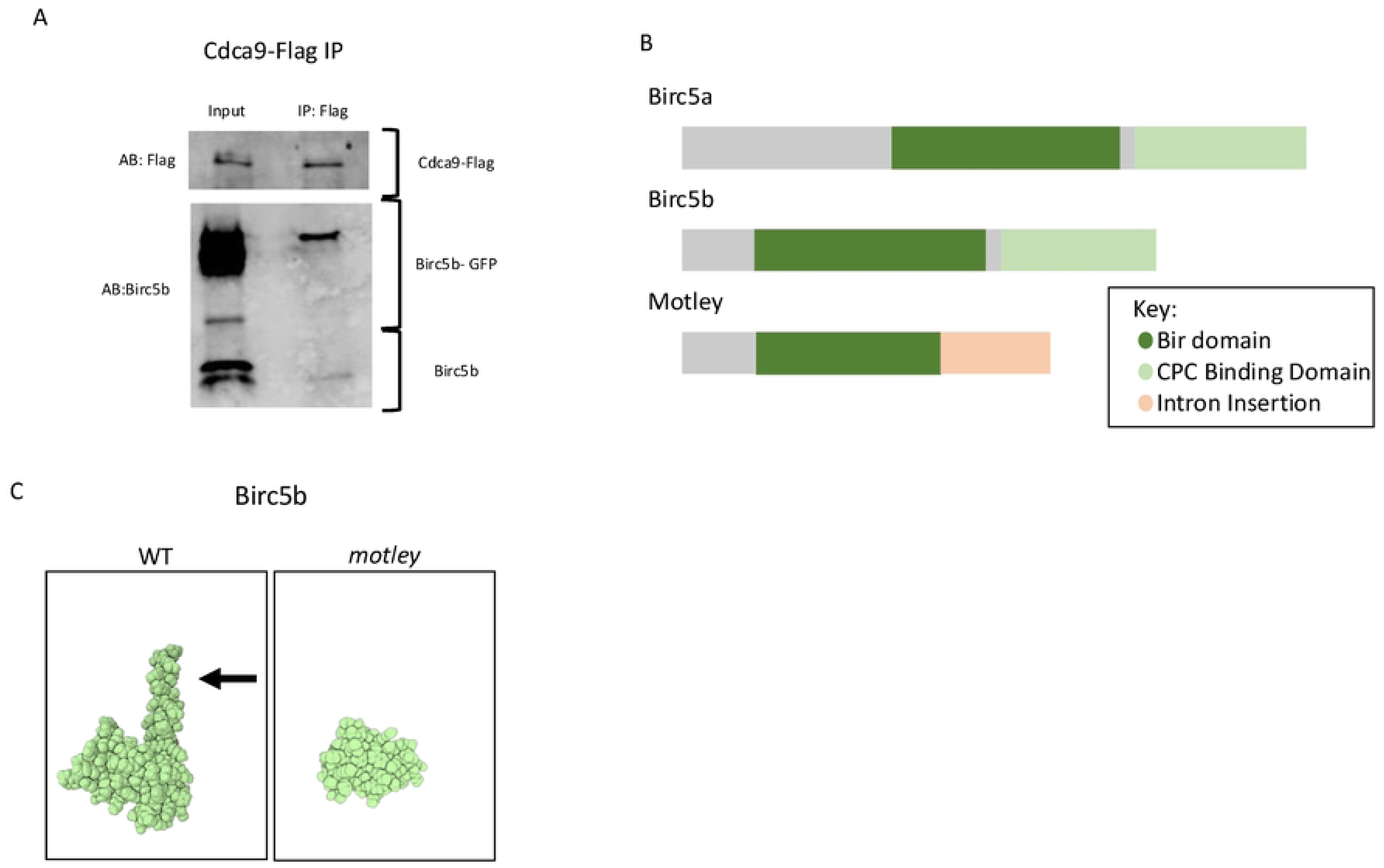
Protein encoded by the *motley/ birc5b* mutant allele cannot interact with the CPC. A) Whole cell lysate from zebrafish embryos injected with *cdca9-flag* and *birc5b-gfp* mRNA subjected to a Co-Immunoprecipitation assay. Cdca9-Flag pulls down both Birc5b-GFP and Birc5b. B) Diagrams highlighting known conserved domains in Birc5a and Birc5b and the predicted truncated product encoded by the *motley (mot^p1aiue^*) mutation(12). The motley mutant protein product lacks the CPC binding domain, preventing it from interacting with Borealin and INCENP via the triple helical bundle. C) A predicted 3-dimensional structure of Birc5b and Motley via Robetta (74) predicts that the truncated product lacks the CPC binding domain (Black arrow).

**Supplementary Figure 4:**
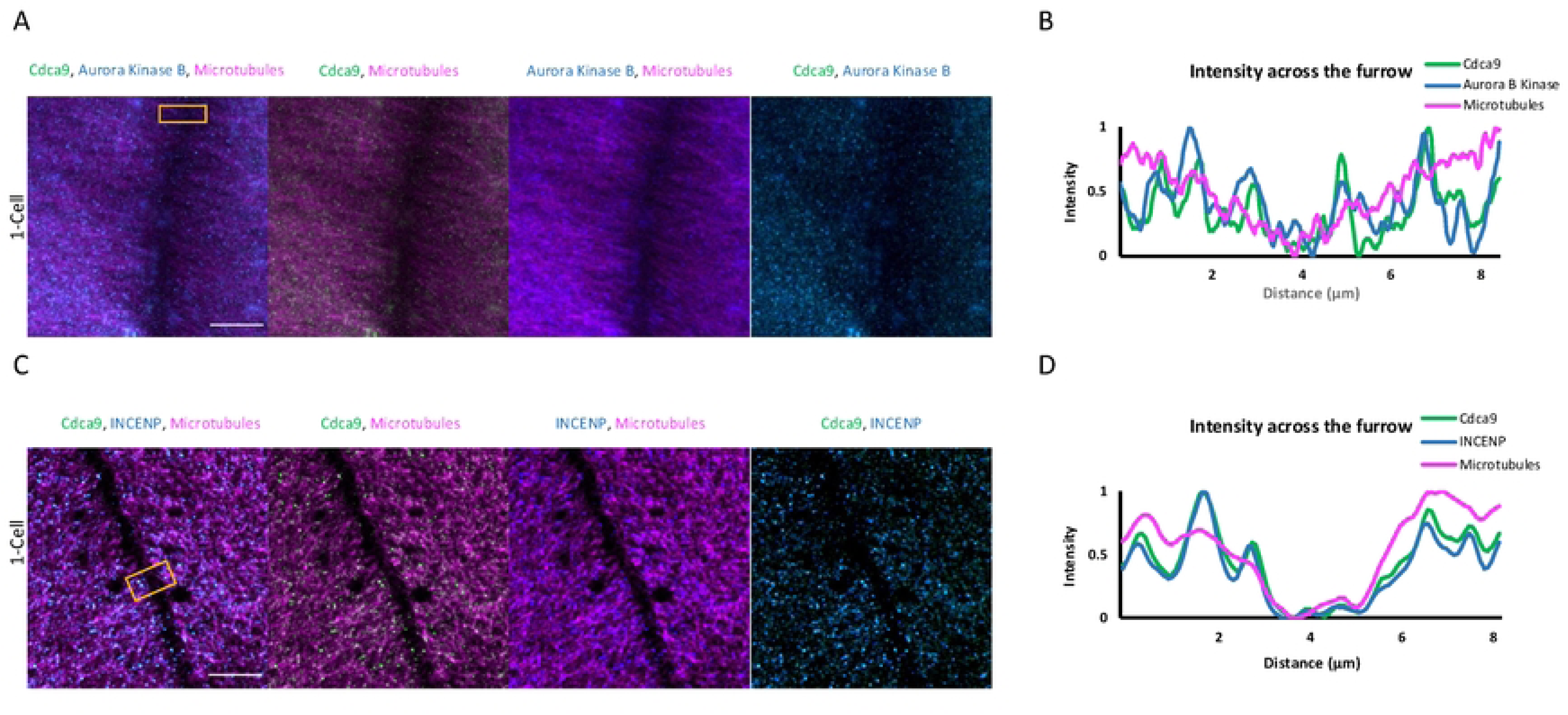
Aurora Kinase B And INCENP are localized to the edge of the furrow. A) Micrographs visualizing the top view (blastodisc) of an early 2-cell embryo labeled for Cdca9, Aurora Kinase B, and microtubules. The general localization pattern shows Cdca9 and Aurora Kinase B localized to the microtubules, not the furrow. B) A Line scan across the furrow (Gold box) shows that the majority of Aurora Kinase B is colocalized with Cdca9. C) Micrographs visualizing the top view (blastodisc) of an early 2-cell embryo labeled for Cdca9, INCENP, and microtubules. The general localization pattern shows Cdca9 and INCENP localized to the microtubules, not the furrow. B) A Line scan across the furrow (gold box) shows that the majority of INCENP colocalizes with Cdca9 and is not found in the furrow. Micrographs are 2D maximum projections of confocal Z-stacks, and the scale bars are 10 μm. Each condition contains 20 ROIs,

**Supplementary Figure 5:**
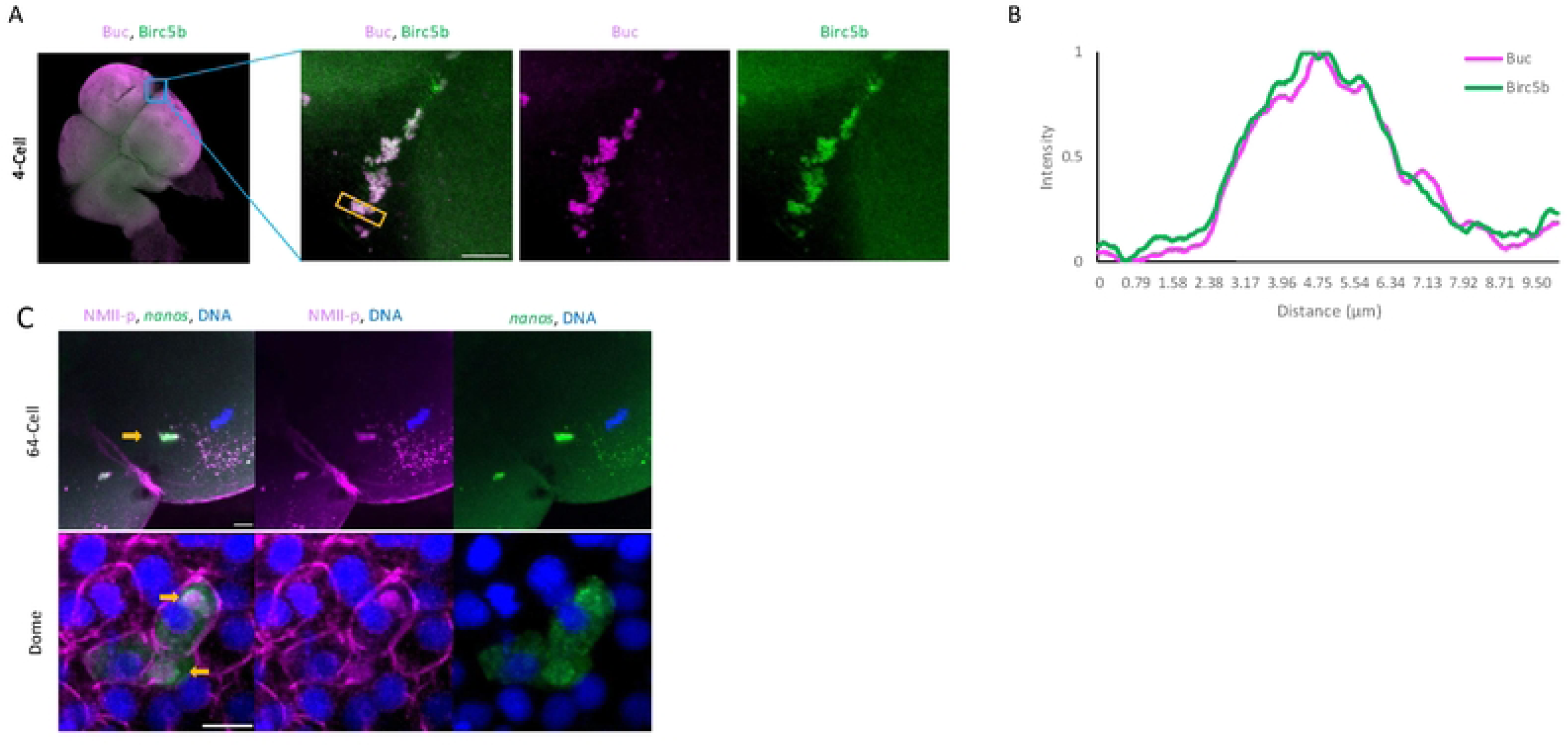
Identification of proteins within the germ plasm mass during early development. A) Micrographs visualizing the top view (blastodisc) of a 4-cell embryo labeled with Birc5b and Buc. The general localization pattern shows Birc5b and Buc are colocalized in the germ plasm mass at the distal end of the furrow. B) A Line scan across the furrow (Gold box) shows that the majority of Birc5b colocalizes with Buc in the germ plasm mass. C) Micrographs of germ plasm localization after it has become cellularized (64-cell stage) and during germ plasm RNA dispersal during the dome stage (∼4.3 hpf). Germ plasm marker *nanos* RNA colocalizes with NMll-p protein at the 64-cell stage. At the dome stage, *nanos* RNA disperses into the cytoplasm of the cell, while NMll-p remains in the germ plasm mass. Micrographs are 2D maximum projections of confocal Z-stacks, and scale bar are 10 μm.

**Supplementary Figure 6:**
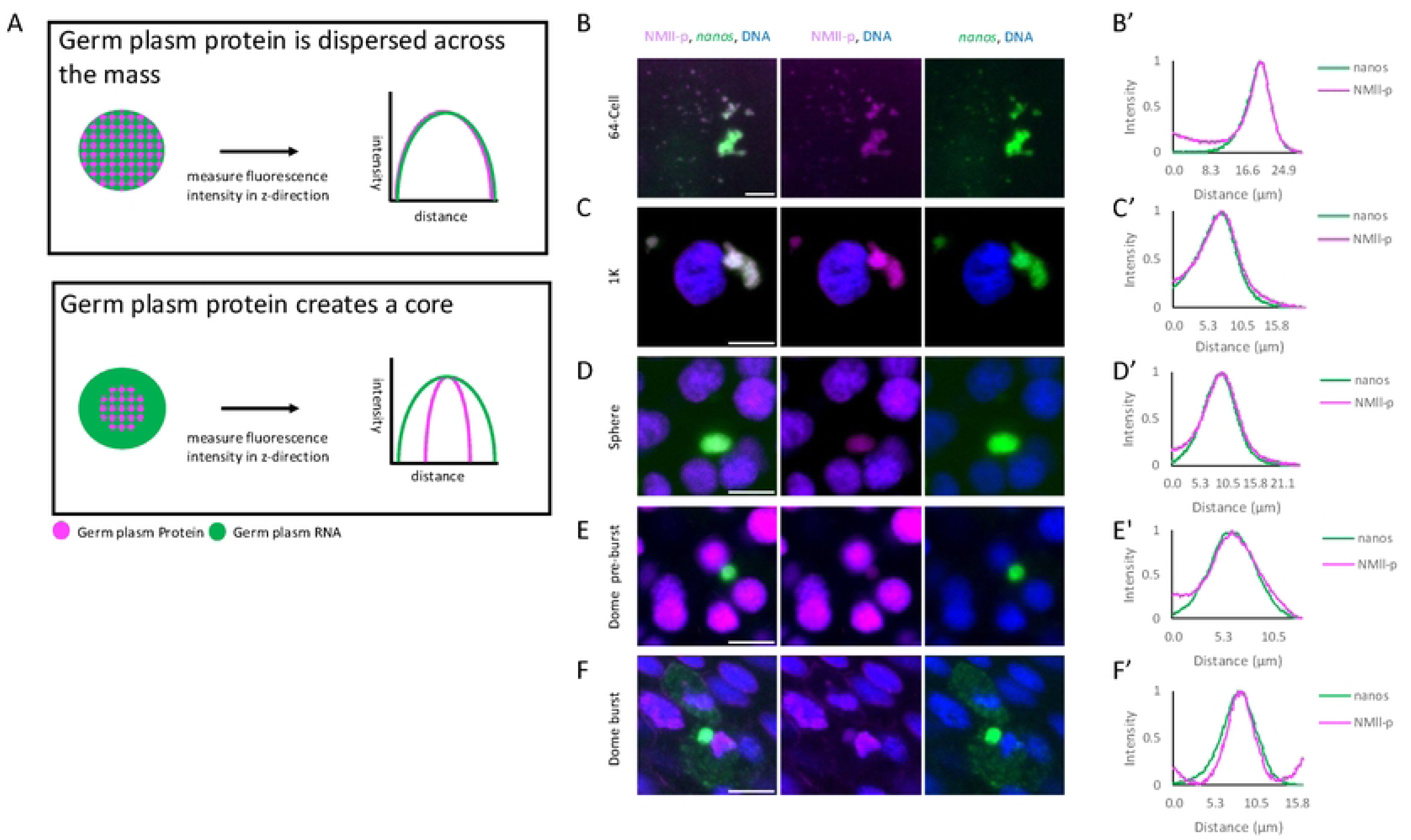
Germ plasm protein is dispersed across the germplasm mass prior to germ plasm RNA dispersal. A) Diagrams showing that the germ protein NMll-p is dispersed across the germ plasm mass or creating a protein core within the germ plasm mass during cleavage stage. B-F) Micrographs of germ plasm within embryonic cells during cleavage (B-D) and dome stage (∼4.3 hpf)(E-F) both before germ plasm RNA dispersal and during germ plasm RNA dispersal. *nanos* RNPs colocalize with NMll-p protein from 64-cell stage to dome stage pre-burst prior to RNP cytoplasmic dispersal. At the dome stage, nanos RNA disperses into the cytoplasm of the cell, while NMll-p remains in the germ plasm mass. B’-F’) Fluorescence intensity profiles of the germ plasm mass showing relative fluorescence intensity in the Z-direction at multiple stages of germ plasm development that are represented by the micrographs. Prior to the dispersal of *nanos* gpRNPs into the cytoplasm (B’-F’), NMll-p is dispersed across the germ plasm mass. All micrographs are 2D maximum projections of confocal Z-stacks, and scale bars are 10 μm.

**Supplementary Figure 7:**
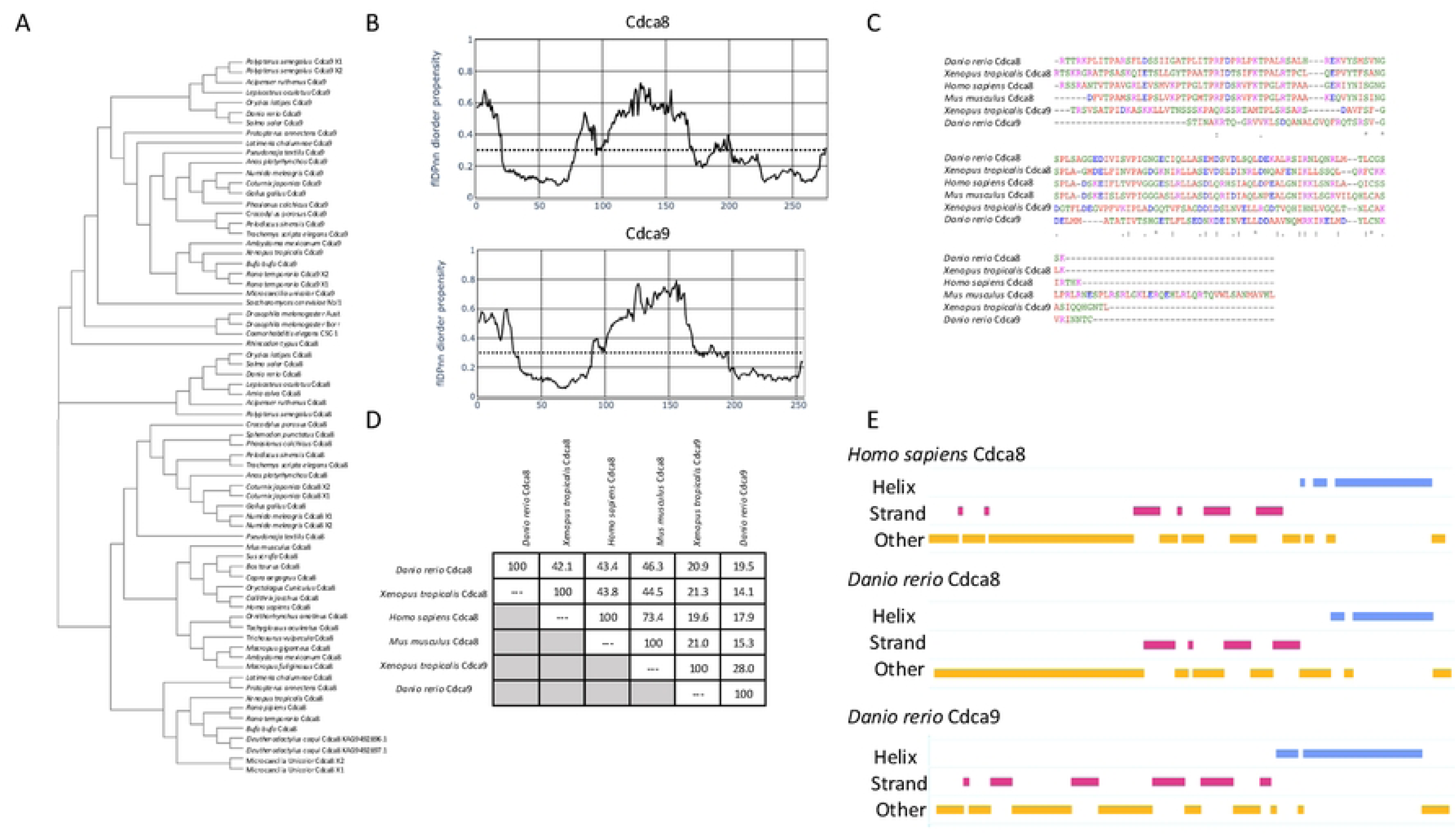
Comparison of Cdca8 and Cdca9 across species. A) Amino Acid phylogenetic tree for Cdca8 and Cdca9 family homologs in selected species. *Danio rerio* Cdca8 clusters with Cdca8 of other species, while Cdca9 clusters with Cdca9 of other species. B) Graph showing the predicted intrinsically disorder regions of zebrafish Cdca8 and Cdca9 C) Multiple sequence alignment comparing the C-terminal of Cdca8 and Cdca9 across species. “*” denotes fully conserved residues, “:” denotes conservation between residues of strongly similar properties, and “.” denotes conservation between residues of weakly similar properties. Amino acids are color-coded according to biochemical properties as follows: red = small, blue = acidic, magenta = basic, green = hydroxyl + sulfhydryl + amine + glycine. The multiple sequence alignments were generated using Clustal Omega (75). D) The percent amino acid identity for the C-terminus of Borealin across species, shows that the tail of Zebrafish Cdca9 is not well conserved. E) Predicted secondary structure of the C-terminal for human Cdca8 and zebrafish paralogs (Cdca8 and Cdca9), shows the conserved alpha helix across species. An increased number of beta strands is observed in zebrafish Cdca9. The secondary structure was predicted using RePROF (76).

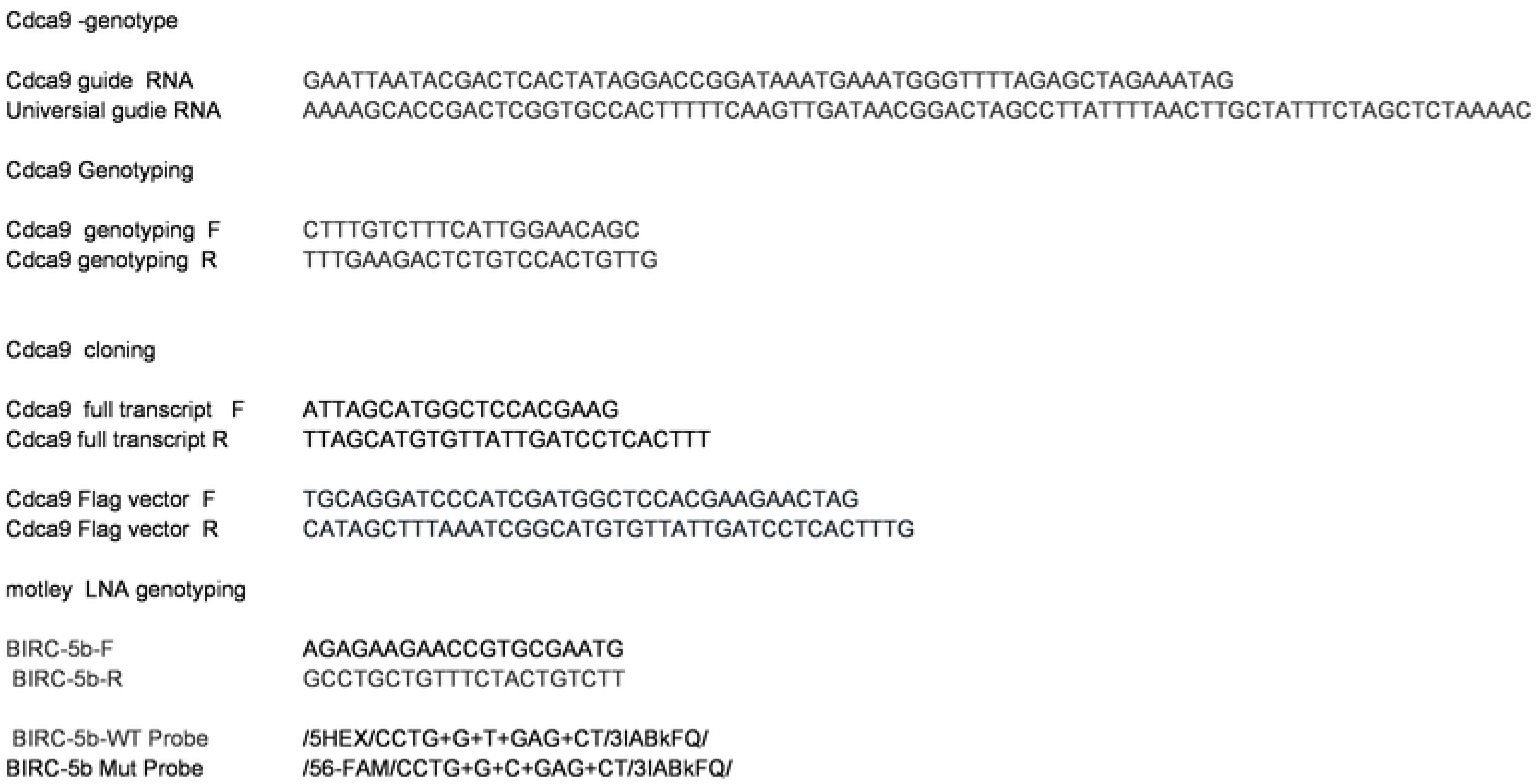

## Notes

### Competing Interest Statement

The authors have declared no competing interest.

